# A p53-Phosphoinositide Signalosome Regulates Nuclear Akt Activation

**DOI:** 10.1101/2021.09.17.460854

**Authors:** Mo Chen, Suyong Choi, Tianmu Wen, Changliang Chen, Narendra Thapa, Vincent L. Cryns, Richard A. Anderson

## Abstract

The tumor suppressor p53 and the phosphoinositide 3-kinase (PI3K)-Akt pathway have fundamental roles in regulating cell growth, apoptosis and are frequently mutated in cancer. Here, we show that genotoxic stress induces nuclear Akt activation by a p53-dependent mechanism that is independent from the canonical membrane-localized PI3K-Akt pathway. Upon genotoxic stress a nuclear p53-PI3,4,5P_3_ complex is generated in regions devoid of membranes by a nuclear PI3K, and this complex recruits all the kinases required to activate Akt and phosphorylate FOXOs, inhibiting DNA damage-induced apoptosis. Wild-type p53 activates nuclear Akt in an on/off fashion upon stress, whereas mutant p53 stimulates high basal Akt activity, indicating a fundamental difference. The nuclear p53-phosphoinositide signalosome is distinct from the canonical membrane-localized pathway and insensitive to PI3K inhibitors currently in the clinic, underscoring its therapeutic relevance.

**In brief:** p53 assembles a PI3K-Akt pathway that regulates nuclear Akt activation independent of the canonical pathway on membranes.

## INTRODUCTION

Phosphoinositide (PI) 3-kinase (PI3K)-mediated generation of PI3,4,5P_3_ downstream of membrane receptors activates the serine/threonine protein kinase Akt, a critical regulator of cell growth, survival, and metabolism, and this pathway is frequently hyperactivated in cancer^1-3^. In the canonical membrane-localized PI3K-Akt pathway, PI3,4,5P_3_ is a required binding lipid that recruits Akt and PDK1 to membranes via their pleckstrin homology (PH) domains^3, 4^ and this enables active PDK1 to phosphorylate T308 on Akt^5^. The subsequent phosphorylation of Akt primarily by mTORC2 on S473 stimulates full Akt activity^3, 6^. Active Akt is present in the cytoplasm^7^ and nucleus^8^. There are reports indicating translocation of active Akt from the cytoplasm to the nucleus^9-11^ and *de novo* nuclear activation of Akt^12-14^. However, the underlying mechanisms of nuclear Akt activation and the functional role of Akt in the nucleus are poorly understood.

The tumor suppressor p53 maintains genome integrity in response to cellular stress and is the most commonly mutated gene in cancer^15, 16^. We have previously demonstrated that p53 interacts with the type I phosphatidylinositol phosphate (PIP) kinase PIPKIα (encoded by *PIP5K1A*), which transfers its product PI4,5P_2_ to the C-terminal domain of p53. The p53-PI4,5P_2_ complex then recruits small heat shock proteins (sHSPs), which bind p53 and stabilize the protein^17^, but the functions of the p53-PI4,5P_2_ complex are unknown. Intriguingly, the nuclear PI3K inositol polyphosphate multikinase (IPMK)^18^ and the 3-phosphatase PTEN^19^ also interact with p53. Moreover, IPMK and PTEN regulate the interconversion of PI4,5P_2_ and PI3,4,5P_3_-bound forms of the nuclear receptor steroidogenic factor 1 (SF-1), which control its transcriptional activity^20, 21^. Additionally, Akt was identified as a potential p53 interactor^22^. These diverse findings suggest that p53 may be systematically targeted by multiple phosphoinositide kinases and phosphatases to produce p53-PI3,4,5P_3_ and regulate Akt activation in the nucleus independently of the canonical membrane-localized pathway.

Here, we demonstrate that genotoxic stress induces nuclear Akt activation by a p53-dependent mechanism. p53-PI4,5P_2_ is converted to p53-PI3,4,5P_3_ in response to cellular stress by the nuclear PI3K inositol polyphosphate multikinase (IPMK), and the 3-phosphatase PTEN reverses this reaction. PI3,4,5P_3_ binding stimulates the assembly of PDK1, mTORC2, Akt, and FOXOs on p53, leading to nuclear activation of Akt and phosphorylation of FOXOs. Inhibition of PIPKIα or IPMK, but not class I PI3Ks, disrupts p53-PI3,4,5P_3_ formation and nuclear Akt activation. The nuclear p53-PI3,4,5P_3_-Akt complex is regulated by cellular stress and attenuates DNA damage, invasion and cell death. Our findings establish a PI3K-Akt pathway that dynamically assembles on p53 to control nuclear Akt activation. This p53-phosphoinositide signalosome is independent of the canonical membrane-localized PI3K-Akt pathway, underscoring its therapeutic relevance.

## RESULTS

### p53 interacts with nuclear Akt and regulates its activation

Although Akt is activated in the nucleus by genotoxic stress and active Akt is observed in the nucleus of cancer cells^8, 12, 13, 23^, the mechanisms of nuclear Akt activation and its function remain enigmatic. To gain mechanistic insights into nuclear Akt activation, we examined the subcellular distribution of phosphorylated Akt under normal and stressed conditions by immunofluorescent (IF) staining. Consistent with prior reports^13, 14^, genotoxic stress increased the nuclear levels of two active Akt phospho-forms (pAkt^T308^ and pAkt^S473^) in the membrane-free nucleoplasm (Fig. 1a,b). Given the critical role of class I PI3K-mediated PI3,4,5P_3_ production in Akt activation in the canonical membrane-localized PI3K-Akt pathway^1, 2^, we postulated a functional role of class I PI3Ks in stress-induced nuclear Akt activation. However, the increase in nuclear pAkt^S473^ levels by cisplatin treatment was not attenuated by the class I PI3K inhibitors alpelisib and buparlisib^24^ at concentrations that suppressed EGF-stimulated Akt activation (Extended Data Fig. 1a-c), indicating that nuclear Akt activation is independent of the canonical PI3K-Akt pathway.

**Figure 1.**
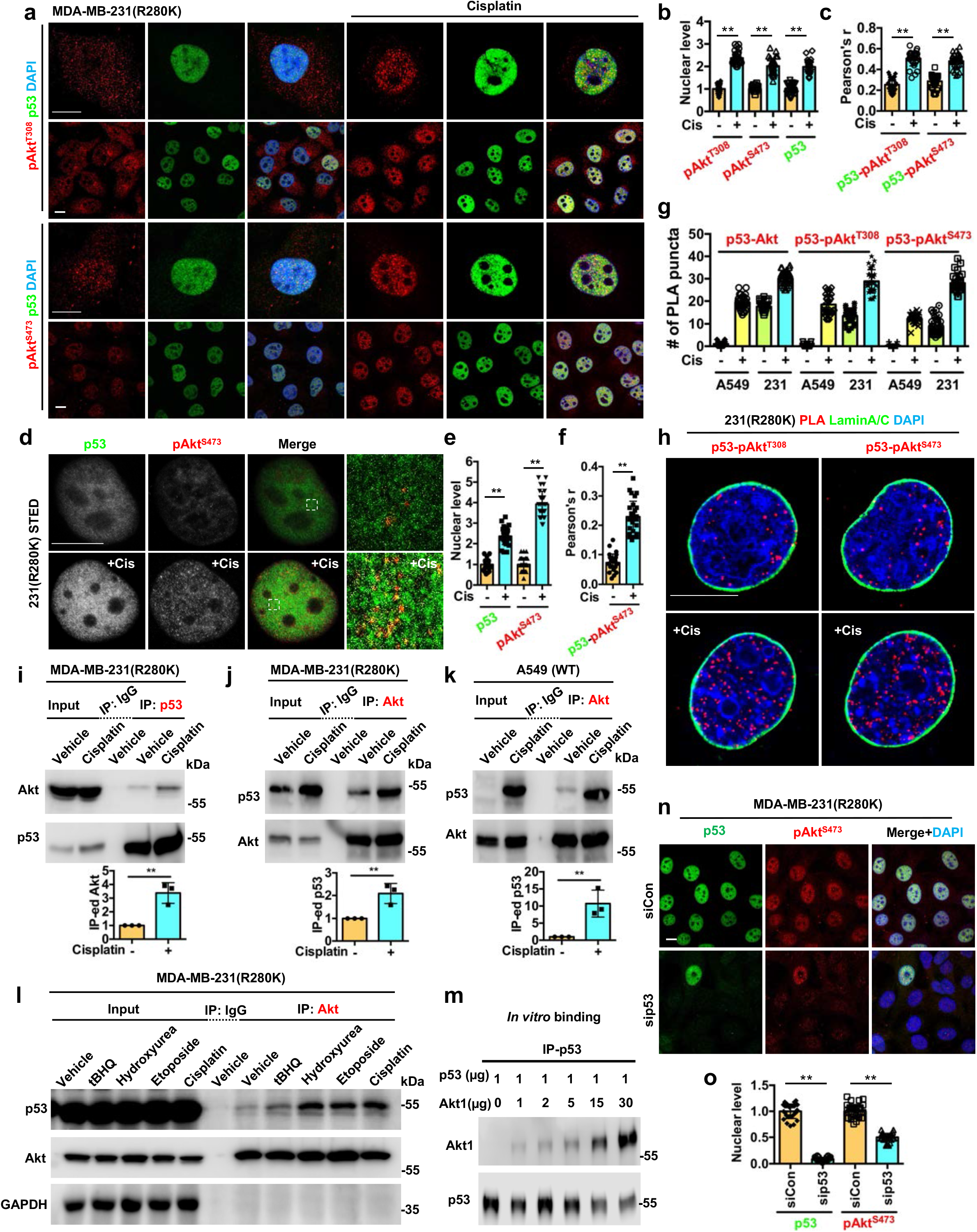
p53 interacts with nuclear Akt and regulates its activation. **a-c**, Confocal images of IF staining against pAkt^T308^/pAkt^S473^ and p53 in MDA-MB-231 cells treated with control vehicle or 30 µM cisplatin for 24 h. The nuclear levels of pAkt^T308^/pAkt^S473^ and p53 (**b**) and their colocalization determined by Pearson’s r (**c**) were quantified. N=30 cells from representative experiments of three repeats. **d-f**, STED super-resolution images of IF staining against pAkt^S473^ and p53 in MDA-MB-231 cells treated with control vehicle or 30 µM cisplatin for 24 h. The nuclear levels of pAkt^S473^ and p53 (**e**) and their colocalization (**f**) were quantified. N=30 cells from representative experiments of three repeats. **g**, Quantification of nuclear PLA complexes of p53-Akt/pAkt^T308^/pAkt^S473^ in A549 and MDA-MB-231 cells treated with control vehicle or 30 µM cisplatin for 24 h. N=30 cells from representative experiments of three repeats. See PLA images in Extended Data Fig. 2a. **h**, PLA of p53-pAkt^T308^/pAkt^S473^ overlaid with nuclear envelope marker Lamin A/C in MDA-MB-231 cells treated with control vehicle or 30 µM cisplatin for 24 h. **i-k**, Immunoprecipitation (IP) of endogenous p53 and Akt from MDA-MB-231 and A549 cells treated with control vehicle or 30 µM cisplatin for 24 h. The Akt IP-ed by mutant p53 (**i**), mutant p53 IP-ed by Akt (**j**), and wild-type p53 IP-ed by Akt (**k**) were analyzed by WB. N=3. **l**, IP of p53 and Akt from MDA-MB-231 cells treated with 100 µM tert-butylhydroquinone (tBHQ), 100 µM hydroxyurea, 100 µM etoposide, 30 µM cisplatin, or control vehicle for 24 h. Representative data of three independent experiments are shown. **m**, *In vitro* binding of recombinant p53 with Akt1. A constant amount of p53 immobilized on anti-p53 agarose was incubated with the indicated increasing amounts of Akt1. p53 was IP’ed and p53-bound Akt1 was analyzed by WB. Representative data of three independent experiments are shown. **n-o**, IF staining of p53 and pAkt^S473^ in MDA-MB-231 cells 48 h after transient transfection with control siRNAs or siRNAs against p53. The nuclear p53 and pAkt^S473^ levels were quantified (**o**). N=30 cells from representative experiments of three repeats. For all panels, data are represented as mean ±SD, p < 0.01 = **, t test. Scale bar: 5 µm.

Akt was identified as a potential p53 interacting protein^22^ and p53 forms a stable nuclear complex with the PIP 5-kinase PIPKIα and the PI3,4,5P_3_ precursor PI4,5P_2_^17^, suggesting a link between nuclear Akt and p53. Consistently, cisplatin increased the colocalization of the active Akt isoforms (pAkt^T308^ and pAkt^S473^) with mutant p53^R280K^ (Fig. 1a-c). Multiple cell stressors augmented nuclear Akt activation and colocalization of pAkt^S473^ with mutant p53 (Extended Data Fig. 1d-g). Stimulated-depletion-emission (STED) microscopy confirmed that genotoxic stress enhanced the colocalization of mutant p53 and pAkt^S473^ in the nucleus, although not all p53 and pAkt^S473^ colocalized (Fig. 1d-f). Genotoxic stress also increased nuclear levels of wild-type 53 and pAkt^S473^and their nuclear colocalization, but there was no stress-induced cytosolic colocalization of p53 and pAkt^S473^ (Extended Data Fig. 1h-n). Cisplatin enhanced the association of both wild-type and mutant p53^R280K^ with total Akt, pAkt^T308^, and pAkt^S473^ in the nucleus as quantified by proximity ligation assay (PLA) (Fig. 1g and Extended Data Fig. 2a). Multiple cellular stressors increased the levels of the p53-pAkt^S473^ complex in the nucleus (Extended Data Fig. 2b). The mutant p53^(R280K)^-pAkt complexes identified by PLA localized to the non-membranous nucleoplasm and were stimulated by genotoxic stress (Fig. 1h). 3D section images and reconstitution confirmed the nuclear localization of the p53-pAkt^S473^complex to membrane-free regions (Extended Data Fig. 2c and Extended Data Videos 1-2). The interactions of both wild-type and mutant p53^R280K^ with Akt were also detected by co-IP and enhanced by cisplatin (Fig. 1i-k and Extended Data Fig. 2d). Multiple stress stimuli augmented mutant p53 binding to Akt (Fig. 1l). Direct binding of recombinant p53 and Akt1 proteins was detected by p53 IP (Fig. 1m). Additionally, the interaction of p53 with Akt1 was quantified by microscale thermophoresis (MST). Akt1 and p53 directly interacted with 10±2 nM affinity (Extended Data Table 1). To determine the functional role of p53 in nuclear Akt activation, we silenced p53 with specific siRNAs and measured nuclear mutant p53^R280K^ and pAkt^S473^ levels. Quantification showed a loss of nuclear pAkt^S473^ upon p53 knockdown (KD) (Fig. 1n,o). Together, these data indicate that p53 interacts with Akt in the nucleus and promotes its activation.

### p53 recruits upstream Akt-activating kinases

To elucidate the mechanisms by which p53 activates nuclear Akt, we examined the association of p53 with the upstream kinases required for cytosolic Akt activation. Consistent with the effects of genotoxic stress on the nuclear localization of Akt, cisplatin increased the nuclear levels of active and nuclear-retained PDK1 (pPDK1^S241^ and pPDK1^S396^, respectively)^25, 26^ and increased their colocalization with mutant p53^R280K^ (Fig. 2a-c). Cisplatin enhanced the association of both wild-type and mutant p53^R280K^ with nuclear PDK1 (total PDK1, pPDK1^S241^, and pPDK1^S396^) and the Sin1 subunit of mTORC2, which contains a PI3,4,5P_3_-specific PH domain^27^, as quantified by PLA (Fig. 2d,e and Extended Data Fig. 3a). The p53-pPDK1 and p53-Sin1 complexes identified by PLA localized to the non-membranous nucleoplasm and were stimulated by genotoxic stress (Fig. 2f). The interaction of both wild-type and mutant p53^R280K^ with PDK1 and Sin1 was also detected by co-IP and stimulated by genotoxic stress (Fig. 2g-j and Extended Data Fig. 2d). Additionally, p53 directly binds to PDK1 and Sin1 with high affinity, 6±0.2 and 20±1 nM, respectively, when quantified by MST (Extended Data Table 1). Collectively, these data establish a robust, stress-induced association of wild-type and mutant p53 with Akt and its activating kinases in the nucleus (Fig. 2k).

**Figure 2.**
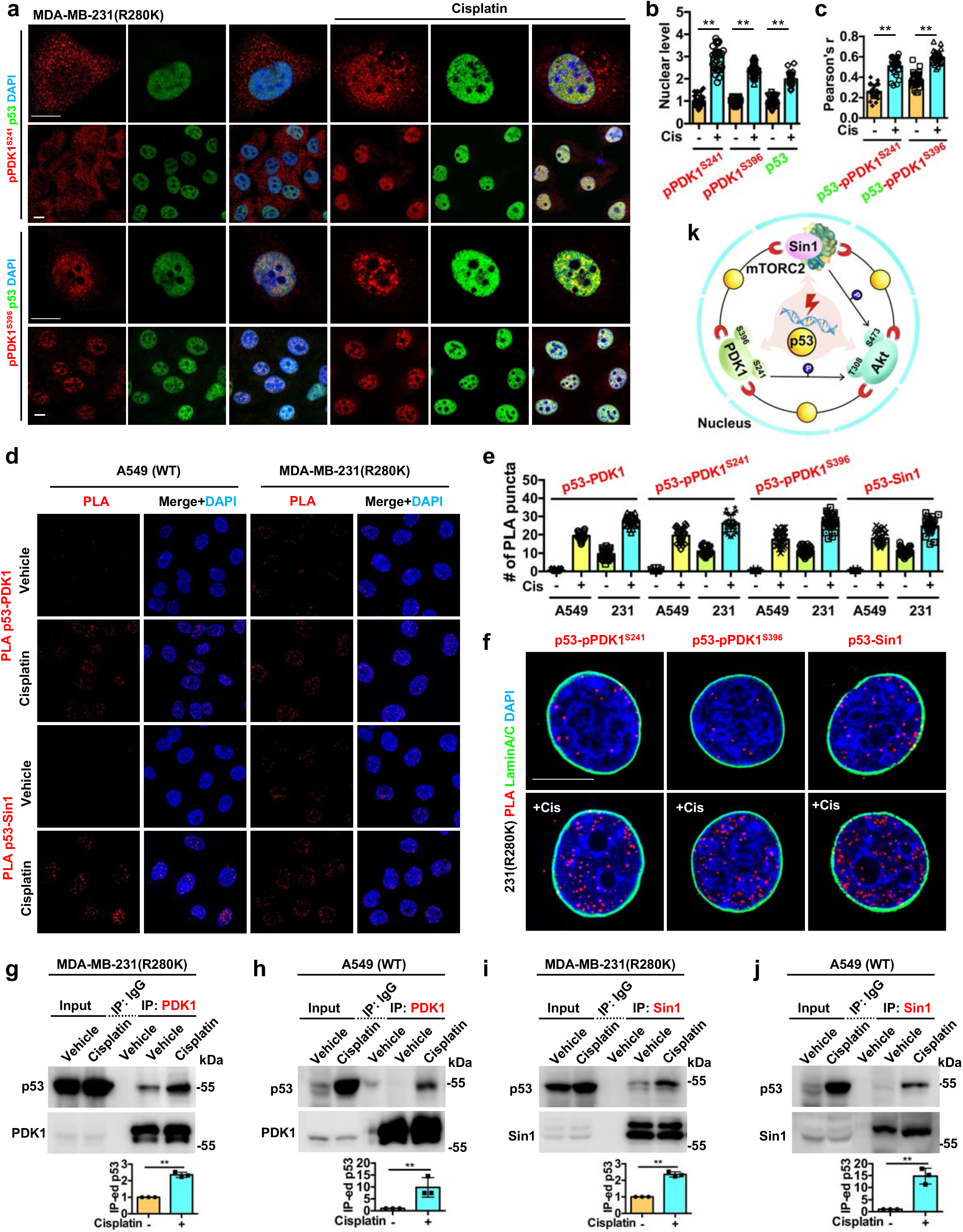
p53 recruits upstream Akt-activating kinases in the nucleus. **a-c**, IF staining against pPDK1^S241^/pPDK1^S396^ and p53 in MDA-MB-231 cells treated with control vehicle or 30 µM cisplatin for 24 h. The nuclear levels of pPDK1^S241^/pPDK1^S396^ and p53 (**b**) and their colocalization (**c**) were quantified. N=30 cells from representative experiments of three repeats. **d-e**, PLA of p53-PDK1/Sin1 in A549 and MDA-MB-231 treated with control vehicle or 30 µM cisplatin for 24 h. The nuclear PLA foci of p53-PDK1/pPDK1^S241^/pPDK1^S396^/Sin1 were quantified (**e**). See images of PLA p53-pPDK1^S241^/pPDK1^S396^ in Extended Data Fig. 3a. **f**, PLA of p53-pPDK1^S241^/pPDK1^S396^/Sin1 overlaid with nuclear envelope marker Lamin A/C in MDA-MB-231 cells treated with control vehicle or 30 µM cisplatin for 24 h. **g-j**, IP of endogenous p53 and PDK1/Sin1 from MDA-MB-231 and A549 cells treated with control vehicle or 30 µM cisplatin for 24 h. The mutant and wild-type p53 IP-ed by PDK1 (**g,h**), and the mutant and wild-type p53 IP-ed by Sin1 (**i,j**) were analyzed by WB. N=3. See also Extended Data Fig. 2d. **k**, Schematic diagram summarizes the nuclear stress-induced p53 complex that activates Akt. For all panels, data are represented as mean ±SD, p < 0.01 = **, t test. Scale bar: 5 µm.

### The nuclear p53-Akt complex targets FOXOs

Akt phosphorylates multiple downstream effectors, including tuberous sclerosis complex 2 (TSC2), glycogen synthase kinase (GSK3), and members of the forkhead box O (FOXO) family^2^. Among these targets, FOXOs are reported to also have direct functional interactions with p53^28-30^. FOXOs directly bind p53^29^ and p53 regulates MDM2 ubiquitination and degradation of FOXOs^30^. The FOXO family, including FOXO1, FOXO3, FOXO4, and FOXO6 in humans, regulate cell senescence and apoptosis in response to genotoxic stress and other stimuli^3, 31^. Akt phosphorylates FOXO proteins at three residues (Fig. 3a), resulting in the nuclear exclusion and degradation of FOXOs to promote cell survival^3^. FOXO1/3, FOXO fragments and phosphorylated forms co-IP’ed with Akt and mutant p53^R280K^ as intact proteins and also proteolytic fragments that were increased by cisplatin (Fig. 3b-d).

**Figure 3.**
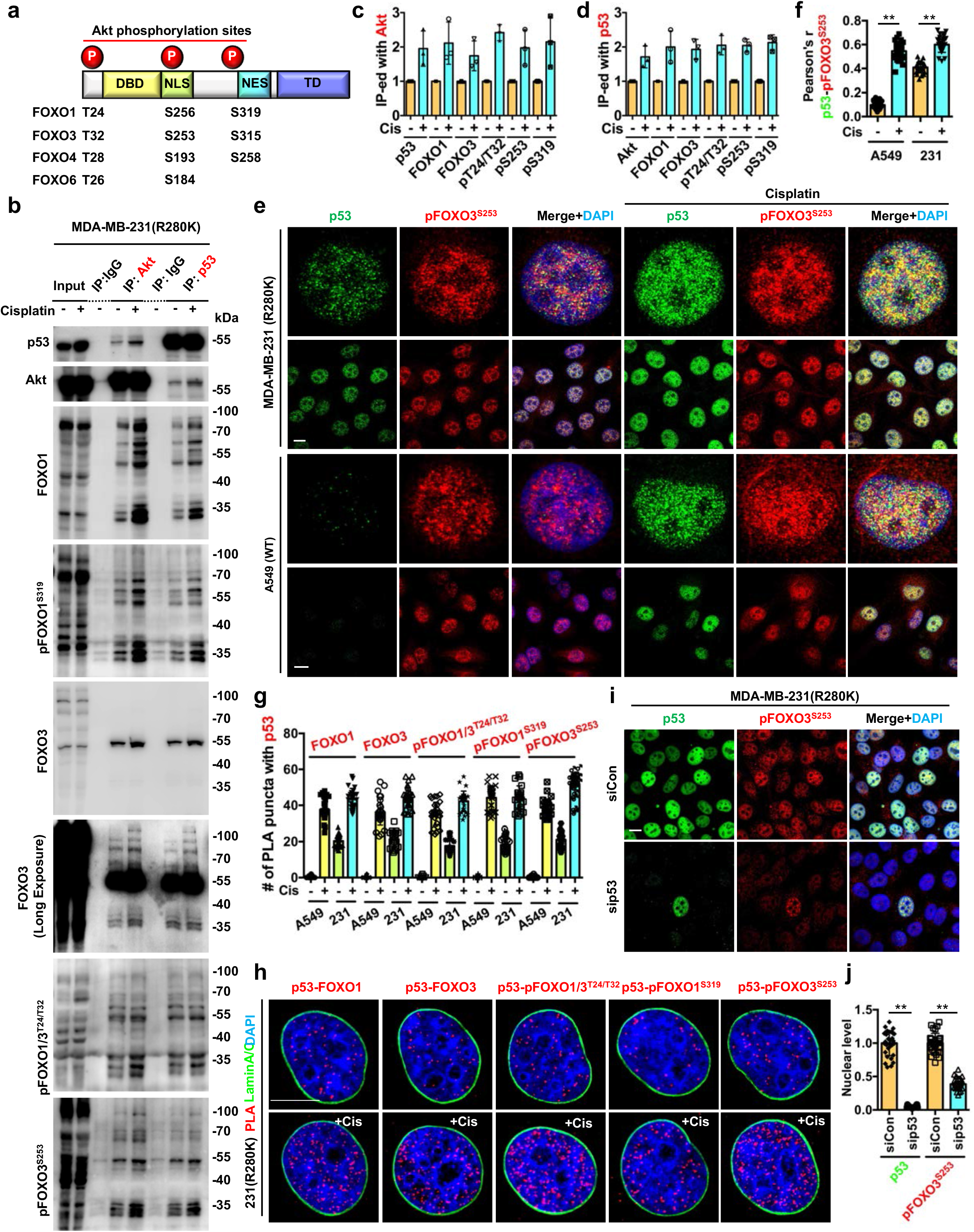
The nuclear p53-Akt complex targets FOXOs. **a**, Domain map of FOXO family members and their phosphorylation sites by Akt. **b-d**, IP analysis of total and phosphorylated forms of FOXOs in association with Akt and p53 in MDA-MB-231 cells treated with control vehicle or 30 µM cisplatin for 24 h. The FOXOs in association with Akt (**c**) and p53 (**d**) were quantified. N=3. **e-f**, IF staining of p53 and pFOXO3^S253^ in A549 and MDA-MB-231 cells treated with control vehicle or 30 µM cisplatin for 24 h. The nuclear colocalization of p53 and pFOXO3^S253^ was quantified (**f**). N=30 cells from representative experiments of three repeats. **g**, Quantification of nuclear PLA complexes of p53 with FOXOs in A549 and MDA-MB-231 treated with control vehicle or 30 µM cisplatin for 24 h. N=30 cells from representative experiments of three repeats. See PLA images in Extended Data Fig. 3b. **h**, PLA of p53-FOXOs overlaid with nuclear envelope marker Lamin A/C in MDA-MB-231 cells treated with control vehicle or 30 µM cisplatin for 24 h. **i-j**, IF staining of p53 and pFOXO3^S253^ in MDA-MB-231 cells 48 h after transient transfection with control siRNAs or siRNAs against p53. The nuclear p53 and pFOXO3^S253^ levels were quantified (**j**). N=30 cells from representative experiments of three repeats. For all panels, data are represented as mean ±SD, p < 0.01 = **, t test. Scale bar: 5 µm.

Wild-type and mutant p53 colocalized with nuclear pFOXO3^S253^ and their colocalization was enhanced by genotoxic stress (Fig. 3e,f). In addition, p53 associated with FOXO1/3 and their phosphorylated forms in the membrane-free nucleoplasm in a stress-regulated manner (Fig. 3g,h and Extended Data Fig. 3b). Functionally, KD of mutant p53 diminished nuclear pFOXO3^S253^ levels (Fig. 3i,j). These findings indicate that the p53-Akt complex recruits and phosphorylates FOXOs in the nucleus, thereby underscoring the functional role of this complex in nuclear Akt and p53 signaling.

### p53 binds and forms a stable complex with PI3,4,5P_3_

Given the essential role of PI3,4,5P_3_ generation and direct binding for Akt activation^1, 2^, we postulated that stress-regulated phosphoinositide linkage on p53 may serve as a nuclear signal to recruit and activate Akt in the nucleus. As we have demonstrated that PI4,5P_2_ forms a stable complex with p53^17^, we determined if PI3,4,5P_3_ also associated with p53. The PI4,5P_2_ and PI3,4,5P_3_ epitopes were detected by fluorescent immunoblotting (IB) of FLAG-immunoprecipitated (IP), ectopically expressed wild-type and mutant p53^175H^, and the interaction of both PI4,5P_2_ and PI3,4,5P_3_ with mutant p53^R175H^ was blocked by the PIPKIα inhibitor ISA-2011B^32^ (Fig. 4a). PI4,5P_2_ and PI3,4,5P_3_ interacted constitutively with diverse mutant p53 proteins and with wild-type p53 in response to cisplatin treatment in all cancer cell lines assayed (Fig. 4b). PI3,4,5P_3_ binding to mutant p53^R280K^ was enhanced by cisplatin treatment and abolished by the PIPKIα inhibitor ISA-2011B^32^, but significantly not by the class I PI3K inhibitors alpelisib and buparlisib (Fig. 4c).

**Figure 4.**
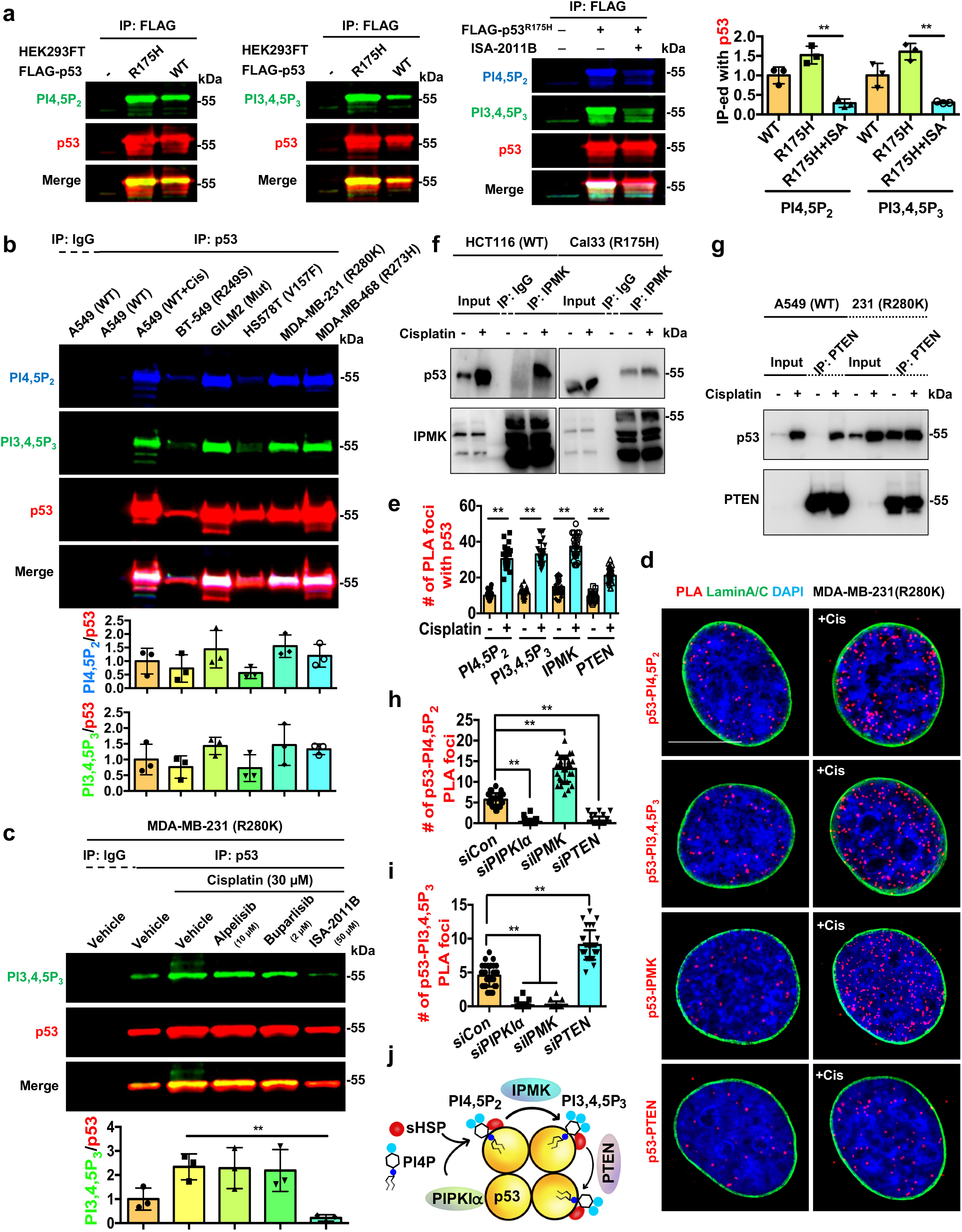
p53 binds PI3,4,5P_3_ and IPMK and PTEN regulate the interconversion of nuclear p53-PI complexes. **a**, Fluorescent IP-WB detects on-site PI4,5P_2_ and PI3,4,5P_3_ association with epitope-tagged p53 downstream of PIPKIα. FLAG-tagged mutant (R175H) or wild-type p53 were transient-transfected into HEK293FT cells for 24 h. Then the cells were treated with 50 µM PIPKIα inhibitor ISA-2011B (ISA) for 24 h. The ectopically-expressed p53 was IP-ed with FLAG antibody and analyzed by WB using fluorescent antibodies detecting PI4,5P_2_, PI3,4,5P_3_, and p53 simultaneously. N=3. **b**, Fluorescent IP-WB detects on-site PI4,5P_2_ and PI3,4,5P_3_ association with endogenous mutant p53 and stress-induced wild-type p53 in a panel of cancer cells. A549 cells expressing wild-type p53 were treated with 30 µM cisplatin or control vehicle for 24 h before being processed for IP against p53. BT-549, GILM2, HS578T, MDA-MB-231, and MDA-MB-468 cells expressing mutant p53 were directly processed for IP against mutant p53. The on-site PI4,5P_2_ and PI3,4,5P_3_ association with p53 was analyzed simultaneously by WB using fluorescent antibodies. N=3. **c**, Fluorescent IP-WB detects stress-induced PI3,4,5P_3_ association with endogenous mutant p53 downstream of PIPKIα but independent from class I PI3Ks. MDA-MB-231 cells were treated with control vehicle or 30 µM cisplatin with or without the presence of the α-specific PI3K inhibitor alpelisib (10 µM), the pan-PI3K inhibitor buparlisib (2 µM), or the PIPKIα inhibitor ISA-2011B (50 µM) for 24 h. Then the cells were processed for IP against p53 and analyzed by WB using fluorescent antibodies detecting PI3,4,5P_3_ and p53 simultaneously. N=3. **d-e**, PLA of p53-PI4,5P_2_/PI3,4,5P_3_/IPMK/PTEN overlaid with nuclear envelope marker Lamin A/C in MDA-MB-231 cells treated with control vehicle or 30 µM cisplatin for 24 h. The nuclear PLA foci were quantified (**e**). N=30 cells from representative experiments of three repeats. **f**, IP of IPMK with wild-type and mutant p53 from HCT116 and Cal33 cells respectively treated with 30 µM cisplatin or control vehicle for 24 h. Representative data of three independent experiments are shown. **g**, IP of PTEN with wild-type and mutant p53 from A549 and MDA-MB-231 cells respectively treated with 30 µM cisplatin or control vehicle for 24 h. Representative data of three independent experiments are shown. **h-i**, Quantification of the nuclear PLA foci of p53-PI4,5P_2_/PI3,4,5P_3_ in MDA-MB-231 cells 48 h after transient transfection with control siRNAs or siRNAs against PIPKIα, IPMK, and PTEN. N=30 cells from representative experiments of three repeats. See PLA images in Extended Data Fig. 5k and knockdown validation by WB in Extended Data Fig. 5l. **j**, Schematic illustration of p53 in a complex with PI4,5P_2_ and PI3,4,5P_3_ downstream of PIPKIα and their interconversion by IPMK and PTEN. For all panels, data are represented as mean ±SD, p < 0.01 = **, t test. Scale bar: 5 µm.

Cisplatin treatment increased nuclear PI3,4,5P_3_ levels and PI3,4,5P_3_ colocalization with mutant p53^R280K^ as determined by immunofluorescence (IF) (Extended Data Fig. 4a-c). PLA revealed that PI4,5P_2_ and PI3,4,5P_3_ associated selectively with cisplatin-activated wild-type p53, while these phosphoinositides are associated constitutively with mutant p53 and these associations were enhanced by multiple cellular stressors (Extended Data Fig. 4d-h). The p53-PI3,4,5P_3_ complexes were abolished by ISA-2011B but not by alpelisib and buparlisib (Extended Data Fig. 4i,j). The p53-PI4,5P_2_ and p53-PI3,4,5P_3_ complexes were localized in the nucleoplasm in regions distinct from nuclear membranes (Fig. 4d,e). 3D section images further confirmed the nuclear localization of the p53-PI3,4,5P_3_ complexes in membrane-free regions (Extended Data Fig. 4k). The interaction of p53 with PI, PI4,5P_2_, and PI3,4,5P_3_ was quantified by MST: PI4,5P_2_ and PI3,4,5P_3,_ but not PI, bound to recombinant p53 (Extended Data Table 1). These data indicate that PI4,5P_2_ and PI3,4,5P_3_ form a stable nuclear complex with stress-activated wild-type and mutant p53 that is dependent on PIPKIα and insensitive to class I PI3K inhibitors.

### IPMK and PTEN regulate the interconversion of nuclear p53-PI complexes

The nuclear PI3K IPMK and the 3-phosphatase PTEN associate with p53^18, 19^ and we confirmed the interactions of IPMK and PTEN with stress-induced wild-type and mutant p53 by co-IP (Fig. 4f,g). IF staining revealed that genotoxic stress enhanced the content of nuclear IPMK and PTEN and their colocalization with p53 (Extended Data Fig. 5a-g). Both IPMK and PTEN associated with stress-activated wild-type p53 and mutant p53^R280K^ in the nucleus in regions distinct from the nuclear membrane as determined by PLA (Fig. 4d,e and Extended Data Fig. 5h-j). IPMK and PTEN direct binding to p53 and this was quantified (Extended Data Table 1). To determine the functional role of PIPKIα, IPMK, and PTEN on p53-PI4,5P_2_/PI3,4,5P_3_ complex formation, we silenced the expression of each enzyme individually with specific siRNAs and then quantified cellular content of mutant p53^R280K^-PI4,5P_2_/PI3,4,5P_3_ complexes by PLA (Fig. 4h,i and Extended Data Fig. 5k,l). PIPKIα knockdown prevented the formation of the p53-PI4,5P_2_ and p53-PI3,4,5P_3_ complexes. IPMK silencing inhibited the formation of p53-PI3,4,5P_3_ complexes while augmenting p53-PI4,5P_2_ complexes. PTEN knockdown increased p53-PI3,4,5P_3_ complexes, consistent with its 3-phosphatase activity, and disrupted p53-PI4,5P_2_ complex formation. These data indicate that IPMK and PTEN dynamically regulate the interconversion of p53-PI4,5P_2_ and p53-PI3,4,5P_3_ complexes downstream of PIPKIα generation of PI4,5P_2_ (Fig. 4j).

### The p53-PI3,4,5P_3_ complex recruits and activates the nuclear Akt pathway

PDK1, Sin1 and Akt are all PI3,4,5P_3_ effectors that require binding to PI3,4,5P_3_ to sustain their activities^27, 33, 34^. These data suggest that the p53-PI3,4,5P_3_ complex serves as a lipid reservoir that supports the Akt pathway activation in the nucleus. In support of this concept, PI3,4,5P_3_ and Akt co-IP’ed with mutant p53^R280K^ and these interactions were enhanced by cisplatin (Fig. 5a,b), indicating p53, PI3,4,5P_3_ and Akt are associated in one complex. KD of p53, PIPKIα or IPMK decreased the interaction of mutant p53 with Akt measured by co-IP under basal or stressed conditions, while PTEN KD enhanced this interaction, underscoring the functional role of PI3,4,5P_3_ in the formation of the p53-Akt complex (Fig. 5c). Furthermore, KD of mutant p53^R280K^, PIPKIα, or IPMK reduced the nuclear pAkt^S473^ and pFOXO3^S253^ content quantified by IF, whereas PTEN KD enhanced nuclear pAkt^S473^ and pFOXO3^S253^ levels (Fig. 5d,e and Extended Data Fig. 6a). Additionally, nuclear mutant p53^R280K^-pAkt^S473^ and p53^R280K^-pFOXO3^S253^ complexes detected by PLA were diminished by KD of p53, PIPKIα, or IPMK, but enhanced by PTEN KD under basal and stressed conditions (Fig. 5f,g and Extended Data Fig. 6b). Notably, the cisplatin-induced nuclear Akt activation was blocked by the PIPKIα inhibitor ISA-2011B, but not by the class I PI3K inhibitors (Fig. 5h,i). These data indicate that PI3,4,5P_3_ bound to p53 recruits and activates Akt in the nucleus and the activity of this nuclear p53-phosphoinositide signalosome is dynamically regulated by phosphatidylinositol kinases and phosphatases and is separate from the canonical PI 3-kinase pathway.

**Figure 5.**
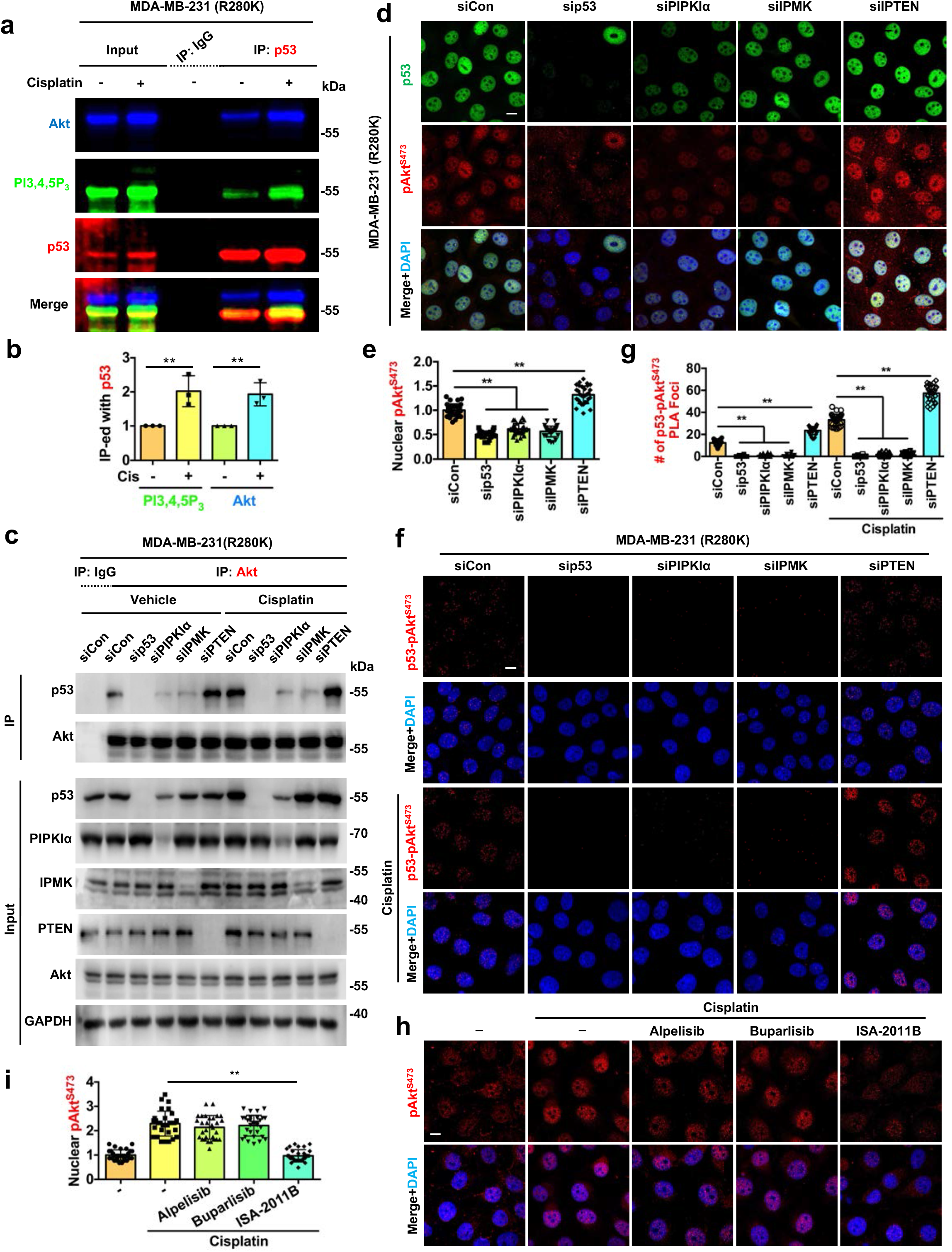
p53-PI3,4,5P_3_ recruits and activates the nuclear Akt pathway. **a-b**, Triple fluorescent IP-WB detects stress-induced Akt association with the p53-PI3,4,5P_3_ complex. MDA-MB-231 cells were treated with 30 µM cisplatin or control vehicle for 24 before being processed for IP against p53 and analyzed by WB using fluorescent antibodies detecting Akt, PI3,4,5P_3,_ and p53 simultaneously. The p53 associated PI3,4,5P_3_ and Akt were quantified (**b**). N=3. **c**, MDA-MB-231 cells were transfected with control siRNAs or siRNAs against p53, PIPKIα, IPMK, and PTEN. 24 h later, cells were treated with 30 µM cisplatin or control vehicle for 24 h before being processed for IP against Akt. The Akt-associated p53 and the input of whole cell lysates were analyzed by WB. Representative images of three independent experiments are shown. **d-e**, IF staining of p53 and pAkt^S473^ in MDA-MB-231 cells 48 h after transient transfection with control siRNAs or siRNAs against p53, PIPKIα, IPMK, and PTEN. The nuclear pAkt^S473^ levels were quantified (**e**). N=30 cells from representative experiments of three repeats. **f-g**, MDA-MB-231 cells were transfected with control siRNAs or siRNAs against p53, PIPKIα, IPMK, and PTEN. 24 h later, cells were treated with 30 µM cisplatin or control vehicle for 24 h before processing for PLA between p53 and pAkt^S473^. The nuclear PLA foci were quantified (**g**). N=30 cells from representative experiments of three repeats. **h-i**, IF staining of pAkt^S473^ in MDA-MB-231 cells treated with control vehicle or 30 µM cisplatin for 24 h with or without the presence of the PI3Kα inhibitor alpelisib (10 µM), the pan-PI3K inhibitor buparlisib (2 µM), or the PIPKIα inhibitor ISA-2011B (50 µM). The nuclear pAkt^S473^ levels were quantified (**i**). N=30 cells from representative experiments of three repeats. For all panels, data are represented as mean ±SD, p < 0.01 = **, t test. Scale bar: 5 µm.

### The p53-PI signalosome links the nuclear PI3K-Akt pathway to DNA-repair and cell survival

PDK1^4, 5, 33^, mTORC2^27^, and Akt^4, 34, 35^ require PI3,4,5P_3_ binding to sustain their activation, and yet the nuclear Akt pathway functions in the non-membranous nucleoplasm. Nuclear pPDK1^S396^ and pFOXO3^S253^ localize to a speckle-like pattern of nuclear PIP-enriched structures^36-38^ (Fig. 2a and 3e). Indeed, double IF staining showed intense and stress-inducible colocalization of pPDK1^S396^ and pFOXO3^S253^ with nuclear PI4,5P_2_ (Fig. 6a,b and Extended Data Fig. 6c). The colocalization of the PDK1-Akt-FOXO complex with the nuclear PI reservoir localizes the nuclear p53-Akt pathway to these non-membranous, PI-enriched domains in the nucleoplasm.

**Figure 6.**
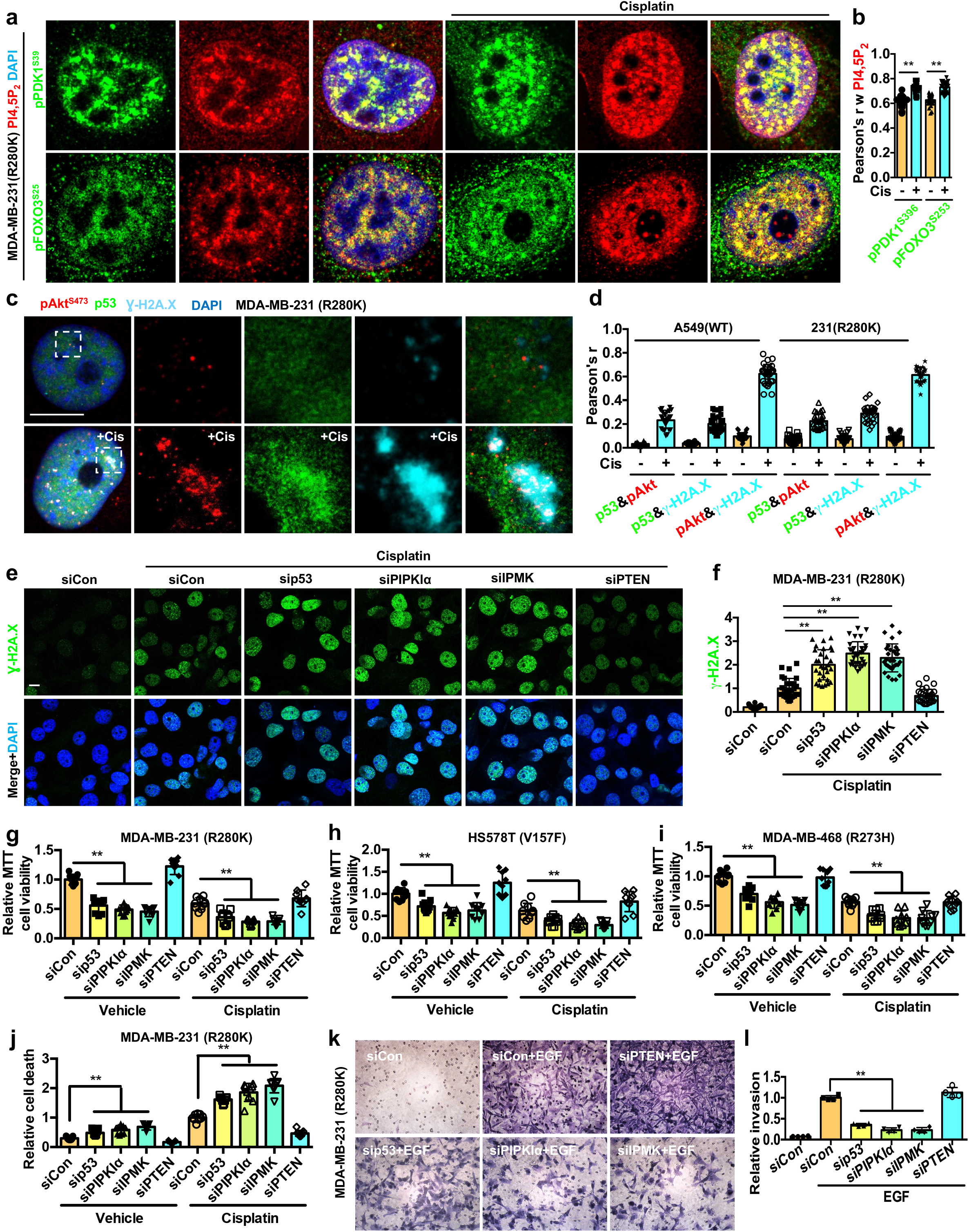
The p53-PI signalosome links the nuclear PI3K-Akt pathway to DNA-repair and cell survival. **a-b**, MDA-MB-231 cells were treated with 30 µM cisplatin or control vehicle for 24 h before being processed for IF staining against PI4,5P_2_ and pPDK1^S396^/pFOXO3^S253^. The nuclear colocalization of PI4,5P_2_ and pPDK1^S396^/pFOXO3^S253^ was quantified (B). N=30 cells from representative experiments of three repeats. Scale bar: 5 µm. See Extended Data Fig. 6c for extended images. **c-d**, STED super-resolution images of IF staining against p53, pAkt^S473^, and the DNA DSB marker Ɣ-H2A.X in MDA-MB-231 or A549 cells treated with 30 µM cisplatin or control vehicle for 24 h. The nuclear colocalization of p53, pAkt^S473^, and Ɣ-H2A.X was quantified (**d**). N=30 cells from representative experiments of three repeats. Scale bar: 5 µm. See Extended Data Fig. 7a for extended images. **e-f**, MDA-MB-231 cells were transfected with control siRNAs or siRNAs against p53, PIPKIα, IPMK, and PTEN. 24 h later, cells were treated with 30 µM cisplatin or control vehicle for 24 h before being processed for IF staining against Ɣ-H2A.X. The nuclear level of Ɣ-H2A.X was quantified (**f**). N=30 cells from representative experiments of three repeats. Scale bar: 5 µm. **g-i**, MDA-MB-231, HS578T, MDA-MB-468 cells were transfected with control siRNAs or siRNAs against p53, PIPKIα, IPMK, and PTEN. 24 h later, cells were treated with 30 µM cisplatin or control vehicle for 24 h before being processed for MTT cell viability assay. N=9 from three independent experiments, each in triplicate. **j**, MDA-MB-231 cells were transfected with control siRNAs or siRNAs against p53, PIPKIα, IPMK, and PTEN. 24 h later, cells were treated with 30 µM cisplatin or control vehicle for 24 h before being processed for cell death detection ELISA. N=9 from three independent experiments, each in triplicate. **k-l**, MDA-MB-231 cells were transfected with control siRNAs or siRNAs against p53, PIPKIα, IPMK, and PTEN. 24 h later, cells were serum-starved for 24 h and then scored for invasion through Laminin-coated transwell inserts with 8 μm pores using a 10 ng/ml EGF chemoattractant for 16 h. The invading cells at the insert bottom were stained with crystal violet, imaged, and quantified based on the extracted dye using a plate reader. N=4. Scale bar: 100 µm. For all panels, data are represented as mean ±SD, p < 0.01 = **, t test.

Our observation that genotoxic stress promotes the assembly of the nuclear p53-PI3,4,5P_3_-Akt-FOXO complex suggested that this complex may play a role in DNA damage repair and/or apoptosis. Indeed, nuclear p53, Akt, FOXOs and phosphoinositides have all been implicated in the DNA damage response^16, 17, 23, 39^ and we have previously localized the p53-PI4,5P_2_ complex to sites of DNA damage^17^. Using the histone H2A.X variant, γ-H2A.X, as a marker for sites of DNA double-strand breaks (DSBs)^40^, STED revealed a specific enrichment of pAkt^S473^ and its colocalization with both wild-type and mutant p53 at DNA damage sites (Fig. 6c,d and Extended Data Fig. 7a). KD of mutant p53, PIPKIα or IPMK increased the cisplatin-induced γ -H2A.X levels, while PTEN silencing diminished γ-H2A.X levels (Fig. 6e,f). Moreover, KD of mutant p53, PIPKIα, or IPMK, sensitized cancer cells to cisplatin treatment and reduced cell invasion, whereas PTEN KD conferred resistance to genotoxic stress and enhanced cell invasion (Fig. 6g-l). Collectively, these results support a pro-survival role of the nuclear p53-PI3,4,5P_3_ complex that recruits and activates Akt at sites of DNA damage to attenuate genotoxic stress.

### The nuclear content of p53 results in Akt activation

Wild-type p53 protein is normally expressed at low levels but is dramatically induced by stress, whereas mutant p53 is expressed at high basal levels due to enhanced protein stability^15, 16^. There are also stochastic variations in the levels of both wild-type and mutant p53 in cells (Fig. 7a-c and Extended Data Fig. 7b-i). To assess the relationship between p53 and active nuclear Akt levels, we quantified the correlation between nuclear p53 and pAkt^S473^. Under basal and stressed conditions, there were strong correlations between nuclear p53 and pAkt^S473^ levels with Pearson’s r greater than 0.7^41^. While the sporadic expression of wild-type p53 segregated the nuclear pAkt levels in an on/off mode, mutant p53 levels correlated with nuclear pAkt expression in a dose-dependent manner (Fig. 7a-c). Moreover, levels of the nuclear Akt target pFOXO3^S253^ also correlated with mutant p53 expression under basal and stressed conditions (Fig. 7d-g). In addition, the nuclear levels of pAkt^S473^ and pFOXO3^S253^ in p53-high cells were significantly higher than those in p53-low cells. To refine this observation, we ectopically expressed Flag-tagged mutant p53^R175H^ in A549 cells and observed that mutant p53 enhanced pAkt^S473^ levels in the nucleus under basal conditions and more so after brief genotoxic stress. The nuclear content of pAkt^S473^ in mutant p53^R175H^ expressing cells was significantly greater than those in mutant p53^R175H^-negative cells (Fig. 7h-k). These results demonstrate that p53 activates nuclear Akt in a dose-dependent fashion and this dose-dependence likely underlies the requirement of cellular stress to stabilize wild-type p53 to assemble the p53-PI3,4,5P_3_-Akt complex and activate nuclear Akt, while hyperstabilized mutant p53 protein ^15, 16^ constitutively activates this pathway with modest augmentation by stress.

**Figure 7.**
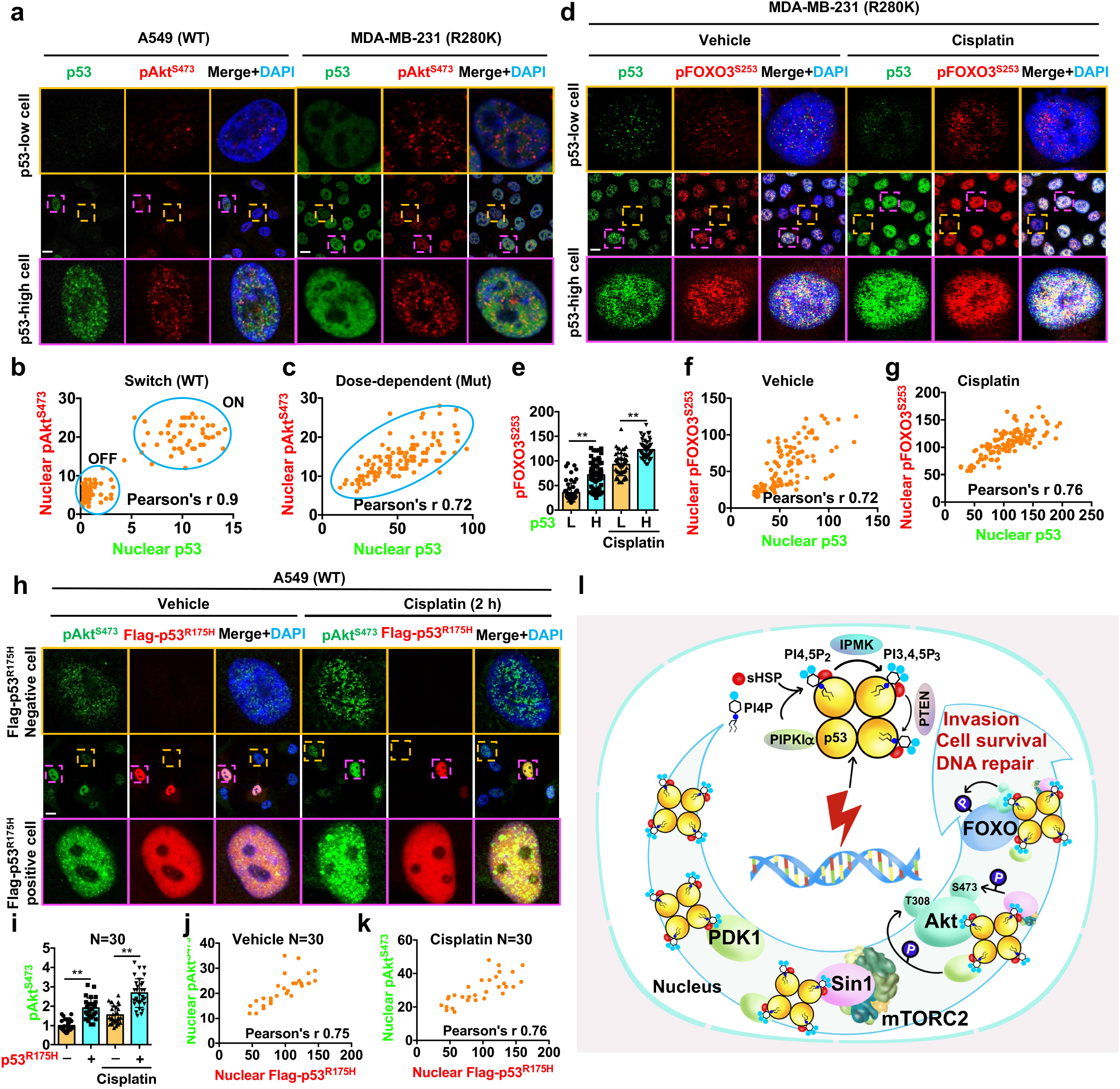
Nuclear p53 levels correlate with Akt activation. **a-c**, A549 and MDA-MB-231 cells after 24 h of treatment with control vehicle were fixed and processed for IF staining for p53 and pAkt^S473^. The nuclear p53 and pAkt^S473^ levels were quantified and the correlation between the nuclear p53 and pAkt^S473^ was calculated by Pearson’s r (**b,c**). N=120 cells from representative experiments of three repeats. See extended images and quantification in Extended Data Fig. 7b-i. **d-g**, MDA-MB-231 cells were treated with 30 µM cisplatin or control vehicle for 24 h and were then fixed and processed for IF staining of p53 and pFOXO3^S253^. The nuclear levels of p53 and pFOXO3^S253^ were quantified. The low (L) and high (H) levels of nuclear p53 were determined based on the average of the p53 levels in the corresponding control vehicle or cisplatin-treated groups (**e**). The correlation between the nuclear p53 and pFOXO3^S253^ was determined by Pearson’s r (**f,g**). N=120 cells from representative experiments of three repeats. **h-k**, A549 cells were transiently transfected with Flag-tagged mutant p53^R175H^ and then 24 h later treated with 30 µM cisplatin or control vehicle for 4 h. The cells were then fixed and processed for IF staining of Flag-tagged mutant p53 and pAkt^S473^. The nuclear pAkt^S473^ levels in Flag-tagged mutant p53^R175H^-negative and -positive cells were quantified (**i**). The correlation between the nuclear levels of Flag-tagged mutant p53^R175H^ and pAkt^S473^ was determined by Pearson’s r (**j,k**). N=30 cells from representative experiments of three repeats. **l**, A schematic model of the stress-induced nuclear p53-phosphoinositide signalosome for Akt activation, DNA-repair, cell survival, and invasion. For all panels, data are represented as mean ±SD, p < 0.01 = **, t test. Scale bar: 5 µm.

## DISCUSSION

Since the discovery of the nuclear generation of PIPs^42^ in regions spatially distinct from known membrane compartments^38^, key components of the canonical membrane-localized cytoplasmic PI3K-Akt pathway have been shown to be present in the nucleus^36^. Nuclear PI signaling enables the synthesis of PIPs and their dynamic interconversions through kinases and phosphatases with no detectable association with membranous structures, implying a distinct nuclear mechanism independent from the cytoplasmic pathway^36, 43^. Accumulating evidence points to nuclear PDK1-mTORC2-Akt activation distinct from the classic membrane-localized cytosolic pathway^8, 44-46^. Notably, the nucleus contains all the machinery necessary for phosphorylating Akt to achieve full activation, including PIP kinases, PI3-kinases, PI4,5P_2_, PI3,4,5P_3_, PDK1, mTORC2, and Akt^8, 36-38^. In the canonical membrane-localized PI3K-Akt pathway, PI3,4,5P_3_ recruits PDK1, mTORC2, and Akt to membranes via their PH domains^27, 33, 34^. Here, we report the discovery of a novel nuclear PI3K-Akt pathway independent of the membrane-localized pathway that is scaffolded on stress-activated wild-type and mutant p53 (Fig. 7l). This nuclear p53-PI signalosome dynamically binds to PIPKIα, IPMK and PTEN to enable stress-regulated modifications of PIPs associated with p53 with exquisite spatial and temporal specificity. The PI3,4,5P_3_ bound to p53 plays an analogous role in the nucleus by recruiting Akt, PDK1, and mTORC2 leading to activation of Akt in the complex.

The nuclear p53-PI3,4,5P_3_-Akt complex also recruits and phosphorylates FOXOs, known Akt substrates implicated in DNA damage-induced apoptosis^23, 39, 47, 48^, and the resulting inactivating phosphorylation of FOXOs appears to mitigate DNA damage and apoptosis, underscoring the cytoprotective function of this complex. Indeed, the localization of the p53-PI3,4,5P_3_-Akt complex at sites of DNA DSBs identified by γ-H2A.X foci supports its role in the DNA damage response.

The discovery of a direct link between p53 and Akt and the dose-dependent induction of nuclear Akt activation by p53 point to a new function of p53. In the absence of stress, wild-type p53 is increased stochastically, activating nuclear Akt in an on/off mode, whereas mutant p53 activates nuclear Akt constitutively, reflecting the hyperstability of mutant relative to the wild-type p53^15, 16^. The loss of transcriptional regulation by mutant p53 and its gain of Akt activation may be underlying mechanisms for its survival and proliferative roles in cancer. Consistent with this idea, knock-in of mutant p53 in mice leads to elevated Akt signaling and tumorigenesis^49^ and expression levels of p53, a surrogate marker for p53 mutational status^50, 51^, correlate with active Akt in human colorectal carcinomas^52^.

The nuclear p53-PI signalosome described here is not only spatially distinct from the membrane-localized PI3K-Akt pathway but also utilizes a different PI3K to generate PI3,4,5P_3_ bound p53, the nuclear PI3K IPMK that interacts with p53^18^. Importantly, the p53-PI signalosome and nuclear Akt activation is resistant to class I PI3K inhibitors including the α-specific inhibitor alpelisib^53^. As such, the discovery of the nuclear p53-PI signalosome has significant therapeutic implications for cancers driven by mutant p53. Finally, the discovery that PIPs tightly associate with key proteins such as p53 and the nuclear poly(A) polymerase Star-PAP^54^ suggests that this may represent a “third messenger” system whereby second messengers in the canonical membrane-localized PI signaling pathway are linked to key regulatory proteins to confer new functional activities.

## ACKNOWLEDGMENTS

We thank Drs. Anjon Audhya, Michael Sussman, and Wolfgang Peti for discussions and comments, and Lance Rodenkirch for technical support. This work was supported in part by a National Institutes of Health grant R35GM134955 (R.A.A.), Department of Defense Breast Cancer Research Program grants W81XWH-17-1-0258 (R.A.A.), W81XWH-17-1-0259 (V.L.C.) and W81XWH-21-1-0129 (V.L.C.), and a grant from the Breast Cancer Research Foundation (V.L.C.).

## AUTHOR CONTRIBUTIONS

MC, SC, TW, CC, NT, VLC, and RAA designed the experiments. MC, SC, TW, CC, and NT performed the experiments. MC, VLC, and RAA wrote the manuscript.

## DECLARATION OF INTERESTS

Authors declare that they have no competing interests.

## METHODS

### Cell culture and constructs

A549, BT-549, Cal33, HCT116, HS578T, MDA-MB-231, MDA-MB-468, SUM1315, HEK293FT, and MCF-10A cells were purchased from ATCC. GILM2 cells were described previously^55^. MCF10A cells were cultured as described^56^ and the other cells were maintained in DMEM (#10-013-CV, Corning) supplemented with 10% fetal bovine serum (#SH30910.03, Hyclone) and 1% penicillin/streptomycin (#15140-122, Gibco). The cell lines used in this study were routinely tested for mycoplasma contamination, and mycoplasma negative cells were used. None of the cell lines used in this study is listed in the database of commonly misidentified cell lines maintained by ICLAC. The p53 constructs used for this work were described previously^17^. Constructs were transfected in mammalian cells by the lipid-based delivery system from Invitrogen (Lipofectamine™3000, #L3000015) according to the manufacturer’s instructions. Typically, 2-5 µg of DNA and 6-10 µl of lipid were used for transfecting cells in 6-well plates. Cells that had at least 80% transfection efficiency were used for further analysis.

### Antibodies and reagents

Monoclonal antibodies against p53 (clone DO-1, #SC-126, Santa Cruz Biotechnology), p53 (clone 7F5, #2527, Cell Signaling), Sin1 (clone D7G1A, #12860, Cell Signaling), Akt (clone C67E7, #4691, Cell Signaling), pAkt^S473^ (clone 193H12, #4058, Cell Signaling), pAkt^T308^(clone D25E6, #13038, Cell Signaling), PTEN (clone 17.A, #MA5-12278, Invitrogen), FOXO1 (clone C29H4, #2880, Cell Signaling), FOXO3 (clone D19A7, #12829, Cell Signaling), GAPDH (clone 0411, #sc-47724, Santa Cruz Biotechnology), and polyclonal antibodies against PIPKIα (PIP5K1A, #9693, Cell Signaling), PDK1 (#3062, Cell Signaling), pPDK1^S241^(#ab131098, Abcam), pPDK1^S396^(#PA5-12888, Invitrogen), pFOXO1^T24^/FOXO3^T32^(#9464, Cell Signaling), pFOXO1^S319^(#2486, Cell Signaling), pFOXO3^S253^(#PA5-37578, Invitrogen) were used in this study. Polyclonal antibodies against PIPKIα□and IPMK were produced as described previously^56^. For conventional immunostaining and PLA analysis of phosphoinositides, anti-PI4,5P_2_ (#Z-P045) and PI3,4,5P_3_ (#Z-P345) antibodies were purchased from Echelon Biosciences. For immunoblotting analyses, antibodies were diluted at a 1:1000 ratio except for p53 (clone DO-1, 1:5000) and GAPDH (clone 0411, 1:5000). For immunoprecipitation, antibody-conjugated agarose was purchased from Santa Cruz Biotechnology, including agarose-conjugated antibodies against p53 (#sc-126AC), PDK1 (#sc-17765AC), Sin1 (#sc-393166AC), Akt (#sc-5298AC), and Flag-tag (#sc-166355AC). For immunostaining analyses and proximity ligation assay (PLA), antibodies were diluted at a 1:100 ratio. Nuclear envelope (Alexa Fluor**®**488 Lamin A/C, clone 4C11, #8617, Cell Signaling, 1:200) and DNA damage site (Alexa Fluor**®**488-γ-H2A.X, clone EP854(2)Y, #ab195188, Abcam, 1:200) markers were used to examine subnuclear regions. For recombinant protein production, the DNA constructs for His-tagged p53, PIPKIα, IPMK, and PTEN were purchased from Genscript and the recombinant proteins were purified as previously described^17^. Recombinant Akt1 (#ab79792) and PDK1 (also known as PDPK1, #ab60834) were purchased from Abcam, and the recombinant Sin1 (also known as MAPKAP1, #TP311745) was purchased from Origene. For the knockdown experiments, pooled-siRNAs targeting the 3’UTR of IPMK (sense 5’-CCAAGAGAGCUGGAAUUCUAUAAUA-3’ and antisense 5’-UAUUAUAGAAUUCCAGCUCUCUUGG-3’; sense 5’-CAAAGGACAACUGUCAGACACAGAA-3’ and antisense 5’-UUCUGUGUCUGACAGUUGUCCUUUG-3’) were purchased from Thermo Fisher Scientific. The ON-TARGETplus siRNA pool against human p53, PIPKIα, and PTEN was purchased from Dharmacon. Non-targeting siRNA (Dharmacon) was used as a control. siRNAs were delivered to cells by RNAiMAX reagent (#13778150, Invitrogen), and knockdown efficiency was determined by immunoblotting. Knockdown efficiency greater than 80% was required to observe phenotypic changes in the study. The PIPKIα inhibitor ISA-2011B (#HY-16937) was purchased from MedChemExpress. Cisplatin (#S1166) and class I PI3K inhibitors alpelisib (BYL719, #S2814) and buparlisib (BKM120, #S2247) were purchased from Sellekchem.

### Immunoprecipitation and immunoblotting

Cells were removed from the medium after the indicated treatment, washed once with ice-cold PBS, and lysed in an ice-cold RIPA lysis buffer system (#sc-24948, Santa Cruz Biotechnology) with 1 mM Na_3_VO_4_, 5 mM NaF, and 1x protease inhibitor cocktail (#11836153001, Roche). The cell lysates were then sonicated at 15% amplitude for 15 s. After sonication, the cell lysates were incubated at 4°C with continuous rotation for 1 h and subsequently centrifuged at maximum speed for 10 min to collect the supernatant. The protein concentration in the supernatant was measured by the Bradford protein assay (#5000201, BIO-RAD) according to the manufacturer’s instructions. Equal amounts of protein were used for further analysis. All antibodies were diluted at a 1:1000 ratio for immunoblotting. For immunoprecipitation, 0.5-1 mg of protein was incubated with 20 µl of antibody-conjugated agarose (Santa Cruz Biotechnology) at 4°C for 24 h. After washing five times with lysis buffer, the protein complex was eluted with SDS sample buffer. For immunoblotting, 5-20 µg of protein were loaded. For immunoblotting of immunoprecipitated complexes, horseradish peroxidase (HRP)-conjugated antibodies were used to avoid non-specific detection of immunoglobulin in the immunoprecipitated samples. HRP-conjugated p53 (#sc-126HRP), PDK1 ((#sc-17765HRP), and Sin1 (#sc-393166HRP) antibodies were purchased from Santa Cruz Biotechnology. Immunoblots were developed by Odyssey Imaging System (LI-COR Biosciences) and the intensity of protein bands was quantified using ImageJ. The unsaturated exposure of immunoblot images was used for quantification with the appropriate loading controls as standards. Statistical analysis of the data was performed with Microsoft Excel, using data from at least three independent experiments.

### Fluorescent IP-WB

Cells were lysed in a RIPA lysis buffer system after the indicated treatment and quantified for protein concentration as described above. For endogenous p53 or FLAG-tagged p53 immunoprecipitation, 0.5-1 mg of cell lysates were incubated with 20 µl anti-p53 (#sc-126 AC, Santa Cruz Biotechnology) or anti-FLAG (#sc-166355AC, Santa Cruz Biotechnology) mouse monoclonal IgG antibody-conjugated agarose at 4°C for 24 h. Normal immunoglobulin (IgG)-conjugated agarose was used as a negative control (#sc-2343, Santa Cruz Biotechnology). After washing 5 times with PBST (PBS with 0.1% Tween 20), the protein complex was eluted with SDS sample buffer. The sample was then boiled at 95°C for 10 min. For immunoblotting, 5-20 µg of protein were loaded. The protein complexes associated with p53 were resolved by SDS-PAGE and transferred onto a PVDF membrane (#IPVH00010, Millipore). The membrane was blocked with 3% BSA in PBS for 1 h at room temperature. For double fluorescent IP-WB detecting p53-PI4,5P_2_/PI3,4,5P_3_ complex, anti-p53 rabbit monoclonal IgG antibody (clone 7F5, #2527, Cell Signaling) at 1:2000 dilution and anti-PI4,5P_2_ mouse monoclonal IgM antibody (#Z-P045, Echelon Biosciences) or PI3,4,5P_3_ mouse monoclonal IgM antibody (#Z-P345, Echelon Biosciences) at 1:2000 dilution were mixed together in blocking buffer with 0.02 % Sodium Azide and incubated with the membrane at 4°C overnight. The next day, the membrane was washed three times with PBST for 10 min each time. For the secondary antibody incubation, goat anti-rabbit IgG antibody conjugated with IRDye 800CW fluorophore (#926-32211, LI-COR) detectable on the 800 nm wavelength channel of the Odyssey Fc Imaging System (LI-COR Biosciences) and goat anti-mouse IgM antibody conjugated with IRDye 680RD fluorophore (#926-68180, LI-COR) detectable on the 700 nm wavelength channel at 1:10000 dilution were mixed together in blocking buffer with 0.01 % SDS and 0.1% Tween 20 and incubated with the membrane at room temperature for 2 h. The membrane was then washed three times with PBST for 10 min each time. The images were subsequently acquired using the 700 and 800 nm wavelength channels simultaneously on the Odyssey Fc Imaging System (LI-COR Biosciences). The p53-associated PI4,5P_2_/PI3,4,5P_3_ complex was visualized by overlapping the 700 and 800 nm channels. For triple fluorescent IP-WB, Alexa fluor 594 Dye (detectable on the 600 nm channel), IRDye 680RD (detectable on the 700 nm channel), and IRDye 800CW (detectable on the 800 nm channel) conjugated antibodies were used together to visualize the complex by overlapping the 600, 700, and 800 nm channels. Statistical analysis of the data was performed with Microsoft Excel, using data from at least three independent experiments.

### *In vitro* binding assay

His-tagged p53 protein was expressed in BL-21(DE3) E. coli (#EC0114, Thermo Fisher), lysed with 1% Brij58, sonicated, and purified with Ni-NTA-agarose (#166038887, Qiagen) as previously described^17^. The eluates were buffer-exchanged into PBS using a dialysis cassette (#66380, Thermo Fisher Scientific), flash-frozen, and stored at -80 °C. Recombinant Akt1 protein was purchased from Abcam (#ab79792). The binding assay was performed in PBST by incubating a constant amount of His-tagged p53 with an increasing amount of Akt1 in the presence of 20 μl anti-p53 antibody-conjugated agarose (#sc-126AC, Santa Cruz). After incubating overnight at 4°C, unbound proteins were removed by washing three times with PBST, and the protein complex was analyzed by immunoblotting.

### Microscale Thermophoresis (MST) Assay

The MST assay was used to measure the binding affinity of purified recombinant proteins *in vitro* as described previously^57^. The target protein was fluorescently labeled by Monolith Protein Labeling Kit RED-NHS 2nd Generation (#MO-L011, Nano Temper) following the manufacturer’s instruction. A sequential titration of unlabeled ligand proteins, PI-PolyPIPosomes, or PI-micelles was made in a Tris-based MST buffer containing 50 mM Tris-HCl, pH 8.0, 50 mM NaCl, 80 mM KCl, and 0.05% Tween-20 and mixed with an equal volume of fluorescently labeled target protein prepared at 10 nM concentration in the same MST buffer, making the final target protein at a constant concentration of 5 nM and the ligand protein or lipid as the gradient. The PI-PolyPIPosomes for MST were purchased from Echelon Biosciences, including PI PolyPIPosomes (#Y-P000), PI4,5P_2_ PolyPIPosomes (#Y-P045), and PI3,4,5P_3_ PolyPIPosomes (#Y-P039). The synthetic PIs for MST were also purchased from Echelon Biosciences, including PI diC16 (#P-0016), PI4,5P_2_ diC16 (#P-4516), and PI3,4,5P_3_ diC16 (#P-3916), which were dissolved in the MST buffer with 5 min sonication to prepare the PI-micelles. The target-ligand mixtures were loaded into Monolith NT.115 Series capillaries (#MO-K022, Nano Temper) and the MST traces were measured by Monolith NT.115 pico, and the binding affinity was auto-generated by MO. Control v1.6 software.

### Immunofluorescence (IF), Confocal and STED Microscopy

For immunofluorescence studies, cells were grown on coverslips coated with 0.2 % gelatin (#G9391, Millipore Sigma). Cells were fixed with 4% paraformaldehyde (PFA) (#sc-281692, Santa Cruz Biotechnology) for 20 min at room temperature followed by washing three times with PBS. Next, the cells were permeabilized with 0.3% Triton-X100 for 10 min and rewashed three times with PBS. The cells were then blocked with 3% BSA in PBS for one hour at room temperature. After blocking, cells were incubated with a primary antibody overnight at 4°C. The cells were then washed three times with PBS and incubated with fluorescent-conjugated secondary antibodies (Molecular Probes) for 1 hour at room temperature. For STED super-resolution microscopy, secondary antibodies conjugated with Abberior® STAR RED/580 dyes were used at 1:200 dilution (#41699 and #52405, Millipore Sigma). After secondary antibody incubation, the cells were washed three times with PBS and nuclei counterstained with 1 µg/ml 4’,6-diamidino-2-phenylindole (DAPI) (#D3571, Invitrogen) in PBS for 30 min at room temperature. The cells were subsequently washed three times with PBS and mounted in Prolong™ Glass Antifade Mountant media (#P36984, Thermo Fisher Scientific). The images were taken by Leica SP8 3xSTED Super-Resolution Microscope, which is both a point scanning confocal and 3xSTED super-resolution microscope. The Leica SP8 3xSTED microscope was controlled by LASX software (Leica Microsystems). All images were acquired using the 100X objective lens (N.A. 1.4 oil). The z-stack images were taken with each frame over a 0.2 µm thickness. For quantification, the mean fluorescent intensity of channels in each cell was measured by LASX. The colocalization of double staining channels was quantified by LASX using Pearson’s correlation coefficient (Pearson’s r), ranging between 1 and −1. A value of 1 represents perfect correlation, 0 means no correlation, and -1 means perfect negative correlation^58^. Pearson’s r greater than 0.7 suggests a strong correlation^41^. The quantitative graph was generated by GraphPad Prism. The images were processed using ImageJ.

### Proximity Ligation Assay (PLA)

PLA was utilized to detect *in situ* protein-protein/PI interaction as previously described^17, 58, 59^. After fixation and permeabilization, cells were blocked before incubation with primary antibodies as in routine IF staining. The cells were then processed for PLA (#DUO92101, Millipore Sigma) according to the manufacturer’s instruction and previously published^58, 59^. The slides post-PLA were further processed for immunofluorescent staining against the nuclear membrane marker (Lamin A/C, #8617, Cell Signaling). The slides were mounted with Duolink® In Situ Mounting Medium with DAPI (#DUO82040, Millipore Sigma). The Leica SP8 confocal microscope detected PLA signals as discrete punctate foci and provided the intracellular localization of the complex. ImageJ was used to quantify the nuclear PLA foci.

### MTT cell viability assay

In 96-well plates, 5×10^4^ cells/well were transfected with control siRNAs or siRNAs targeting p53, PIPKIα, IPMK, and PTEN for 48 h. The cells were then treated with control vehicle or 30 μM cisplatin for 20 h. Next, the cells were incubated with 100 μl of fresh medium plus 10 μl of the 12 mM MTT stock solution from the Vybrant® MTT cell proliferation assay kit (#V13154, Thermo Fisher Scientific) in the presence of the control vehicle or 30 μM cisplatin. After incubation for 4 hours at 37°C, all but 25 μl of the medium was removed from each well and 50 μl of DMSO was added. The mixture was incubated at 37°C for 10 min, and the absorbance was read at 540 nm using a Synergy HTX Multi-Mode Microplate reader (BioTek Instruments, Inc.).

### Cell death detection ELISA

In 96-well plates, 5×10^4^ cells/well were transfected with control siRNAs or siRNAs targeting p53, PIPKIα, IPMK, and PTEN for 48 h. The cells were then treated with control vehicle or 30 μM cisplatin for 24 h. Next, the cells were lysed and processed for cell death detection ELISA assay by following the manufacturer’s instruction (#11774425001, Millipore Sigma). At the end of the assay, the absorbance was read at 405 nm using a Synergy HTX Multi-Mode Microplate reader.

### Transwell Invasion Assay

The bottom polycarbonate filter surface of a 6.5 mm diameter insert with 8 μm pores in a 24-well plate (#3422, Corning) was coated with 10 μg/ml of Laminin (#CC095, Millipore) diluted in PBS for 3 h at 37°C. MDA-MB-231 cells were transiently transfected with control siRNAs or siRNAs targeting p53, PIPKIα, IPMK, and PTEN, then serum-starved for 24 h. Next, 5×10^4^ transfected cells were suspended in 200 μl serum-free medium containing 0.5% BSA and then were plated in the upper insert chamber in 500 μl serum-free medium with 0.5% BSA. 10 ng/ml EGF was added to the lower chamber. Cells were allowed to invade for 16 h at 37°C. Cells on the bottom of the filter were then fixed with 4% PFA and stained with 0.1% Crystal Violet. The stained cells were imaged using an EVOS M5000 microscope (Thermo Fisher Scientific). At the end of the assay, the dye was extracted from the cells and quantified by measuring the optical density at 570 nm using a Synergy HTX Multi-Mode Microplate reader.

### Statistics and Reproducibility

Two-tailed unpaired *t*-tests were used for pair-wise significance unless otherwise indicated. We note that no power calculations were used. Sample sizes were determined based on previously published experiments where significant differences were observed ^17^. Each experiment was repeated at least three times independently, and the number of repeats is defined in each figure legend. We used at least three independent experiments or biologically independent samples for statistical analysis.

### Resource and Data Availability

All data supporting the findings of this study are available from the corresponding authors on reasonable request.

## SUPPLEMENTAL INFORMATION

**Extended Data Fig. 1 p53 associates with stress-induced active Akt in the nucleus a**, WB analysis of pAkt^S473^ in MDA-MB-231 cells after 24 h starvation and treatment with 10 µM PI3Kα inhibitor alpelisib, 2 µM pan-PI3K inhibitor buparlisib, or control vehicle, followed by 5 min stimulation with 50 ng/ml EGF. N=3.

**b-c**, IF staining of pAkt^S473^ in MDA-MB-231 cells treated with control vehicle or 30 µM cisplatin for 24 h with or without the presence of the PI3Kα inhibitor alpelisib (10 µM) or the pan-PI3K inhibitor buparlisib (2 µM). The nuclear pAkt^S473^ levels were quantified (**c**). N=30 cells from representative experiments of three repeats.

**d-g**, IF staining of p53 and pAkt^S473^ in MDA-MB-231 cells treated with 100 µM tBHQ, 100 µM hydroxyurea, 100 µM etoposide, 30 µM cisplatin, or control vehicle for 24 h. The nuclear levels of pAkt^S473^ (**e**) and p53 (**f**) and their colocalization (**g**) were quantified. N=30 cells from representative experiments of three repeats.

**h-k**, IF staining of p53 and pAkt^S473^ in A549 and MDA-MB-231 cells treated with 30 µM cisplatin or control vehicle for 24 h. The nuclear levels of p53 and pAkt^S473^ (**j**) and their colocalization (**k**) were quantified. N=30 cells from representative experiments of three repeats.

**l-n**, Two-panel section of IF staining against p53 and pAkt^S473^ in A549 cells treated with 30 µM cisplatin or control vehicle for 24 h. The middle section focusing on the nucleus and the bottom section focusing on the cytosol were demonstrated in the model (**l**). The nuclear and cytosolic colocalization of p53 and pAkt^S473^ was quantified (**n**). N=30 cells from representative experiments of three repeats.

For all panels, data are represented as mean ±SD, p < 0.01 = **, t test. Scale bar: 5 µm.

**Extended Data Fig. 2 p53 associates with the stress-induced nuclear Akt pathway a**, PLA of p53-Akt/pAkt^T308^/pAkt^S473^ in A549 and MDA-MB-231 cells treated with control vehicle or 30 µM cisplatin for 24 h. See quantification in Fig. 1g.

**b**, PLA of p53-pAkt^S473^ in MDA-MB-231 cells treated with 100 µM tBHQ, 100 µM hydroxyurea, 100 µM etoposide, 30 µM cisplatin, or control vehicle for 24 h. The nuclear PLA foci were quantified. N=30 cells from representative experiments of three repeats.

**c**, 3D section of PLA between p53 and pAkt^S473^ overlaid with Lamin A/C in MDA-MB-231 cells treated with control vehicle or 30 µM cisplatin for 24 h. Each frame of the 3D sections was over a 0.2 μm thickness.

**d**, IP of p53 and Akt pathway components from MDA-MB-231 cells treated with 30 µM cisplatin or control vehicle for 24 h. N=3.

For all panels, data are represented as mean ±SD, p < 0.01 = **, t test. Scale bar: 5 µm.

**Extended Data Fig. 3 p53 associates with PDK1 and FOXOs in the nucleus**

**a**, PLA of p53-pPDK1^S241^/pPDK1^S396^ in A549 and MDA-MB-231 cells treated with control vehicle or 30 µM cisplatin for 24 h. See quantification in Fig. 2e.

**b**, PLA of p53-FOXOs in A549 and MDA-MB-231 cells treated with vehicle or 30 µM cisplatin for 24 h. See quantification in Fig. 3g.

For all panels, scale bar: 5 µm.

**Extended Data Fig. 4 p53 associates with PI4,5P**_**2**_ **and PI3,4,5P**_**3**_ **in the nucleus**

**a-c**, IF staining against p53 and PI3,4,5P_3_ in MDA-MB-231 cells treated with 30 µM cisplatin or control vehicle for 24 h. The nuclear level of PI3,4,5P_3_ (**b**) and its colocalization with p53 (**c**) were quantified. N=30 cells from representative experiments of three repeats.

**d-e**, PLA of p53-PI4,5P_2_/PI3,4,5P_3_ in A549, MCF-10A, MDA-MB-231, and SUM1315 cells treated with 30 µM cisplatin or control vehicle for 24 h. The nuclear PLA foci were quantified (**e**). N=30 cells from representative experiments of three repeats.

**f-h**, PLA of p53-PI4,5P_2_/PI3,4,5P_3_ in MDA-MB-231 cells treated with 100 µM tBHQ, 100 µM hydroxyurea, 100 µM etoposide, 30 µM cisplatin, or control vehicle for 24 h. The nuclear PLA foci were quantified (**g,h**). N=30 cells from representative experiments of three repeats.

**i-j**, PLA of p53-PI3,4,5P_3_ in MDA-MB-231 cells treated with control vehicle or 30 µM cisplatin with or without the presence of the PI3Kα inhibitor alpelisib (10 µM), the pan-PI3K inhibitor buparlisib (2 µM), or the PIPKIα inhibitor ISA-2011B (50 µM) for 24 h. The nuclear PLA foci were quantified (**j**). N=30 cells from representative experiments of three repeats.

**k**, 3D section of PLA between p53 and PI3,4,5P_3_ overlaid with Lamin A/C in MDA-MB-231 cells treated with control vehicle or 30 µM cisplatin treatment for 24 h. Each frame of the 3D sections was over a 0.2 μm thickness.

For all panels, data are represented as mean ±SD, p < 0.01 = **, t test. Scale bar: 5 µm.

**Extended Data Fig. 5 IPMK and PTEN associate with p53 and mediate p53-bound PI4,5P**_**2**_ **and PI3,4,5P**_**3**_ **interconversion in the nucleus**

**a-c**, IF staining against p53 and IPMK in MDA-MB-231 cells treated with 30 µM cisplatin or control vehicle for 24 h. The nuclear level of IPMK (**b**) and its colocalization with p53 (**c**) were quantified. N=30 cells from representative experiments of three repeats.

**d-g**, IF staining against p53 and PTEN in A549 or MDA-MB-231 cells treated with 30 µM cisplatin or control vehicle for 24 h. The nuclear level of PTEN (**f**) and its colocalization with p53 (**g**) were quantified. N=30 cells from representative experiments of three repeats.

**h-j**, PLA of p53-IPMK/PTEN in A549 and MDA-MB-231 cells treated with control vehicle or 30 µM cisplatin for 24 h. The nuclear PLA foci were quantified (**i,j**). N=30 cells from representative experiments of three repeats.

**k-l**, PLA of p53-PI4,5P_2_/PI3,4,5P_3_ in MDA-MB-231 cells 48 h after of transient transfection with control siRNAs or siRNAs against PIPKIα, IPMK, and PTEN (**k**). See quantification in Fig. 4h,i. The knockdown efficiency was validated by WB (**l**).

For all panels, data are represented as mean ±SD, p < 0.01 = **, t test. Scale bar: 5 µm.

**Extended Data Fig. 6 Nuclear FOXO and PDK1 phosphorylation are mediated by the p53-PI signalosome**

**a**, IF staining of p53 and pFOXO3^S253^ in MDA-MB-231 cells 48 h after transient transfection with control siRNAs or siRNAs against p53, PIPKIα, IPMK, and PTEN. The nuclear pFOXO3^S253^ levels were quantified. N=30 cells from representative experiments of three repeats.

**b**, MDA-MB-231 cells were transfected with control siRNAs or siRNAs against p53, PIPKIα, IPMK, and PTEN. 24 h later, cells were treated with 30 µM cisplatin or control vehicle for 24 h before being processed for PLA between p53 and pFOXO3^S253^. The nuclear PLA foci were quantified. N=30 cells from representative experiments of three repeats.

**c**, IF staining against PI4,5P_2_ and pPDK1^S396^/pFOXO3^S253^ in MDA-MB-231 cells treated with 30 µM cisplatin or control vehicle for 24 h. See Fig. 6a,b for the nuclear region and quantification of the nuclear colocalization of PI4,5P_2_ and pPDK1^S396^/pFOXO3^S253^.

For all panels, data are represented as mean ±SD, p < 0.01 = **, t test. Scale bar: 5 µm.

**Extended Data Fig. 7 p53 and active Akt target to DNA damage sites in the nucleus**

**a**, STED super-resolution images of IF staining against p53, pAkt^S473^, and the DNA DSB marker Ɣ-H2A.X in MDA-MB-231 and A549 cells treated with 30 µM cisplatin or control vehicle for 24 h. See quantification in Fig. 6d.

**b-e**, A549 cells were treated with 30 µM cisplatin or control vehicle for 24 h and then were fixed and processed for IF staining of p53 and pAkt^S473^. The nuclei were counter-stained by DAPI. The nuclear levels of p53 and pAkt^S473^ were quantified. The low (L) and high (H) levels of nuclear p53 were determined based on the average p53 levels in the corresponding control vehicle- or cisplatin-treated groups (**c**). The correlation between nuclear p53 and pAkt^S473^ was determined by Pearson’s r (**d,e**). N=120 cells from representative experiments of three repeats.

**f-i**, MDA-MB-231 cells were treated with 30 µM cisplatin or control vehicle for 24 h and then fixed and processed for IF staining against p53 and pAkt^S473^. The nuclei were counter-stained by DAPI. The nuclear levels of p53 and pAkt^S473^ were quantified. The low (L) and high (H) levels of nuclear p53 were determined based on the average of the p53 levels in the corresponding control vehicle or cisplatin-treated groups (**g**). The correlation between the nuclear p53 and pAkt^S473^ was determined by Pearson’s r (**h,i**). N=120 cells from representative experiments of three repeats.

For all panels, data are represented as mean ±SD, p < 0.01 = **, t test. Scale bar: 5 µm.

**Extended Data Table 1**

**The binding affinity of the nuclear p53-PI signalosome components with p53**

The interaction of recombinant fluorescently-labeled p53 and non-labeled signalosome components was quantitated by microscale thermophoresis (MST) assay. A constant concentration of fluorescently-labeled p53 (target, 5 nM) was incubated with increasing concentrations of non-labeled ligands to calculate the binding affinity. The binding affinity determined by MST are shown as indicated K_d_ values. MST was performed using a Monolith NT.115 pico, and the binding affinity was auto-generated by MO. Control v1.6 software. Data are represented as mean±SD; n=3.

**Extended Data Video 1**

**3D reconstitution of the p53-pAkt**^**S473**^ **complex in the nucleoplasm of MDA-MB-231 cells under basal conditions**

MDA-MB-231 cells were treated with control vehicle for 24 h before being processed for PLA between p53 and pAkt^S473^ (Red). Lamin A/C stained the nuclear envelopes (Green), and the nuclei were counter-stained by DAPI (Blue). The z-stack images were taken with a Leica SP8 confocal microscope with each frame over a 0.2 μm thickness. The video of the 3D reconstitution of a representative cell from three independent experiments was generated by LASX.

**Extended Data Video 2**

**3D reconstitution of the p53-pAkt**^**S473**^ **complex in the nucleoplasm of MDA-MB-231 cells under stress**

MDA-MB-231 cells were treated with 30 µM cisplatin for 24 h before being processed for PLA between p53 and pAkt^S473^ (Red). Lamin A/C stained the nuclear envelopes (Green), and the nuclei were counter-stained by DAPI (Blue). The z-stack images were taken with a Leica SP8 confocal microscope with each frame over a 0.2 μm thickness. The video of the 3D reconstitution of a representative cell from three independent experiments was generated by LASX.

## Notes

### Competing Interest Statement

The authors have declared no competing interest.

