## Extended Data Figures for "A p53-Phosphoinositide Signalosome Regulates Nuclear Akt Activation"

#### Extended Data Fig. 1

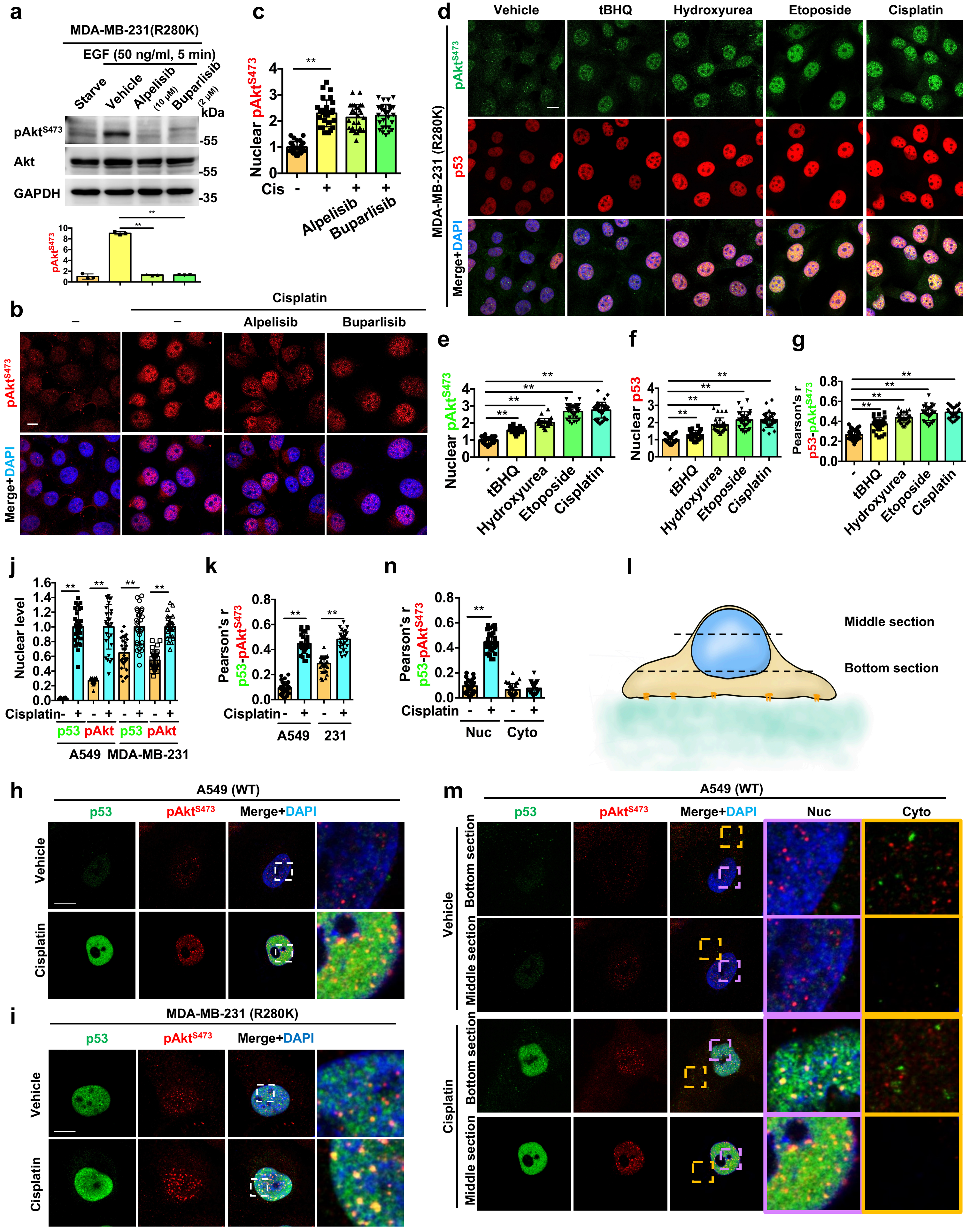

### Extended Data Fig. 2

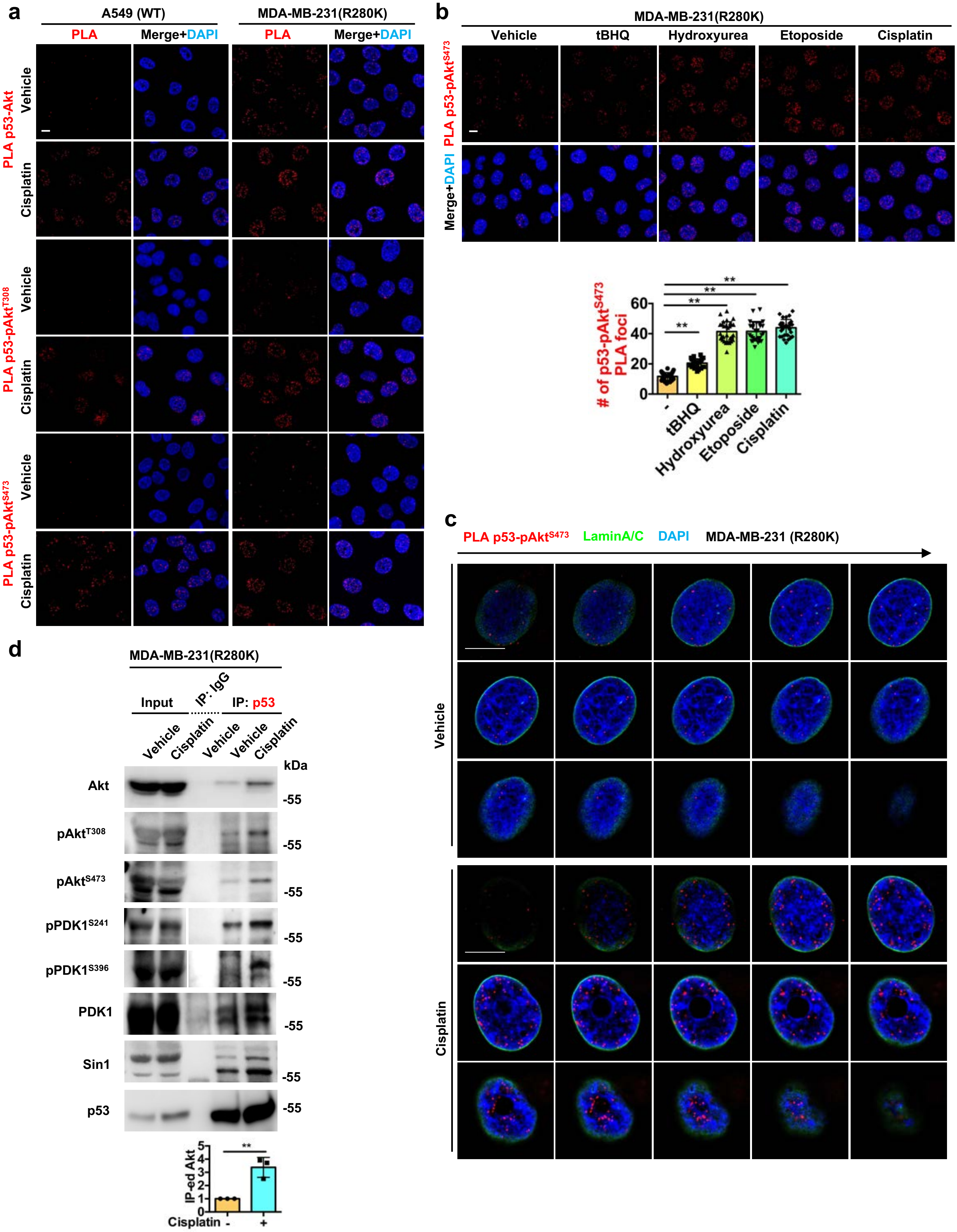

#### Extended Data Fig. 3

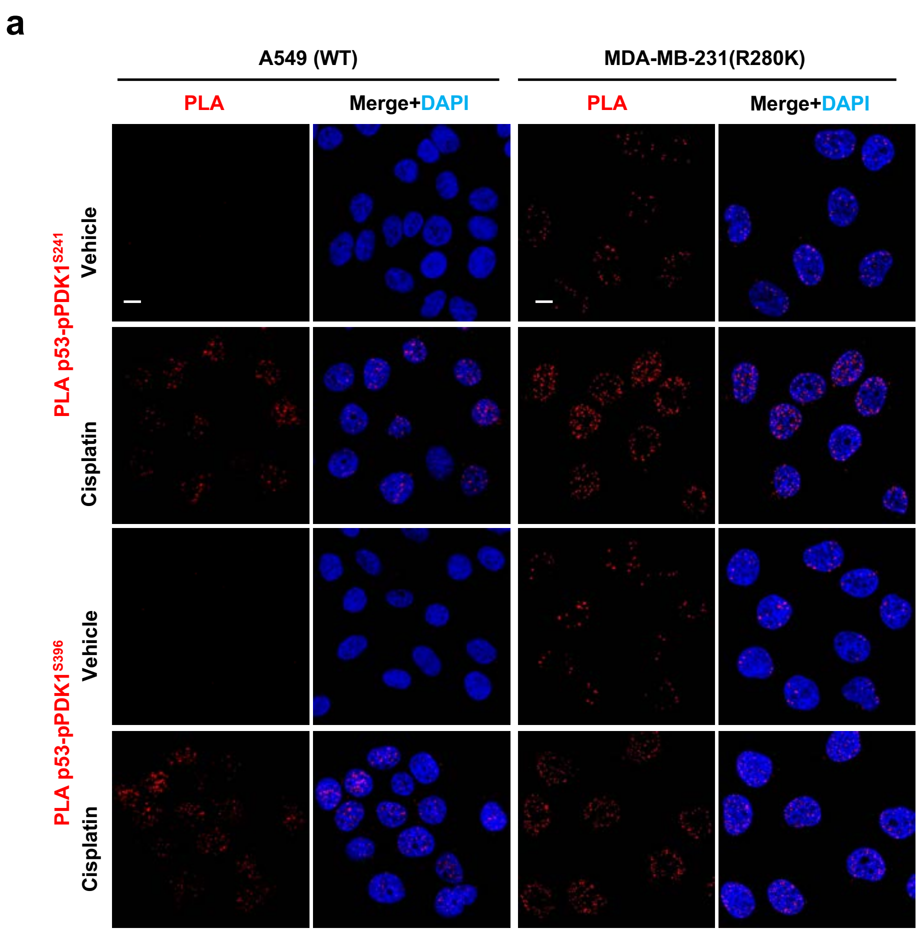


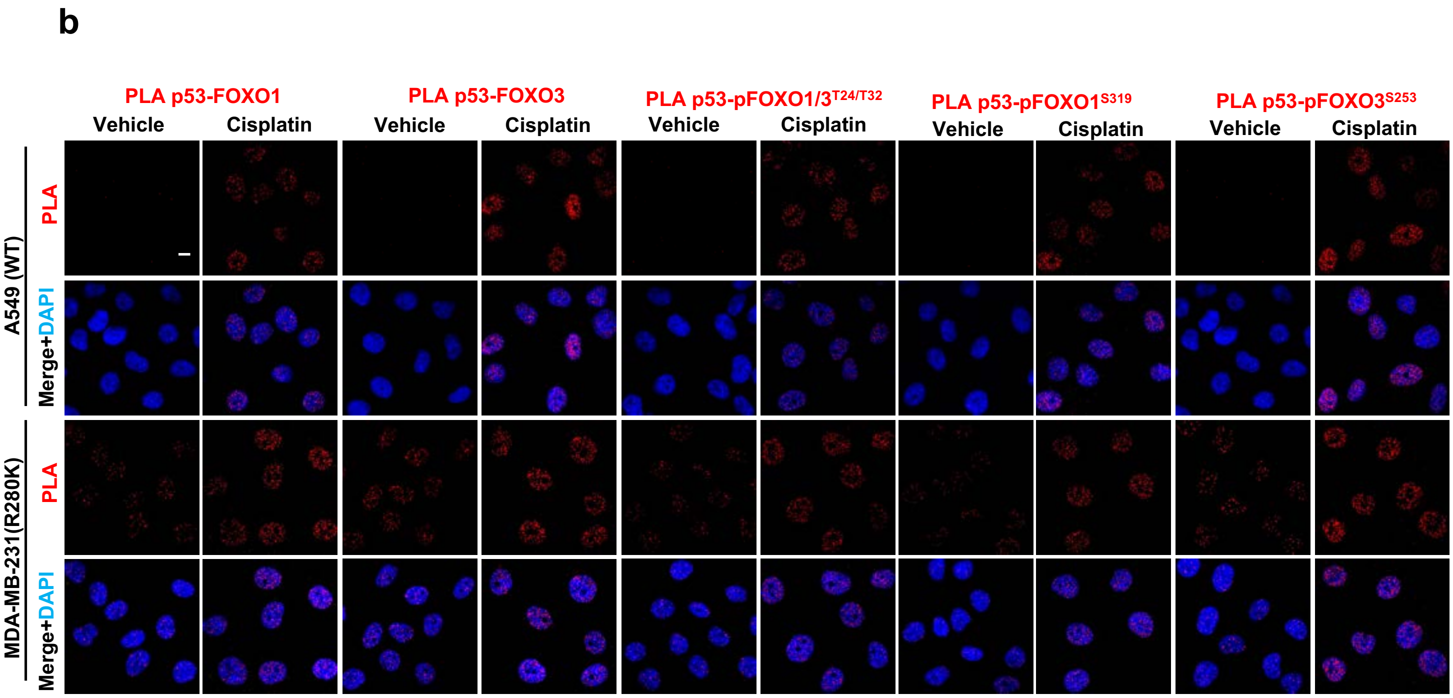

### Extended Data Fig. 4

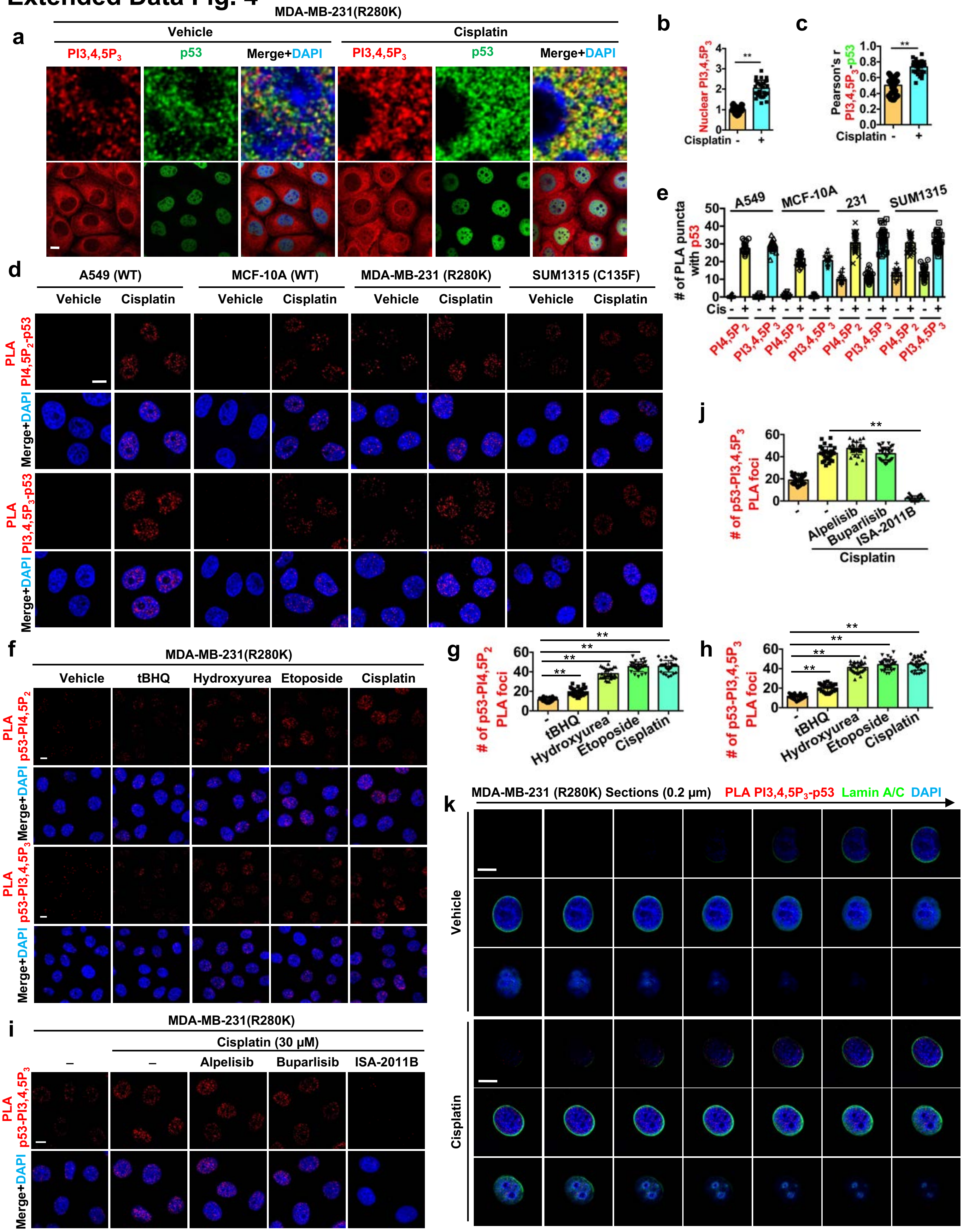

### Extended Data Fig. 5

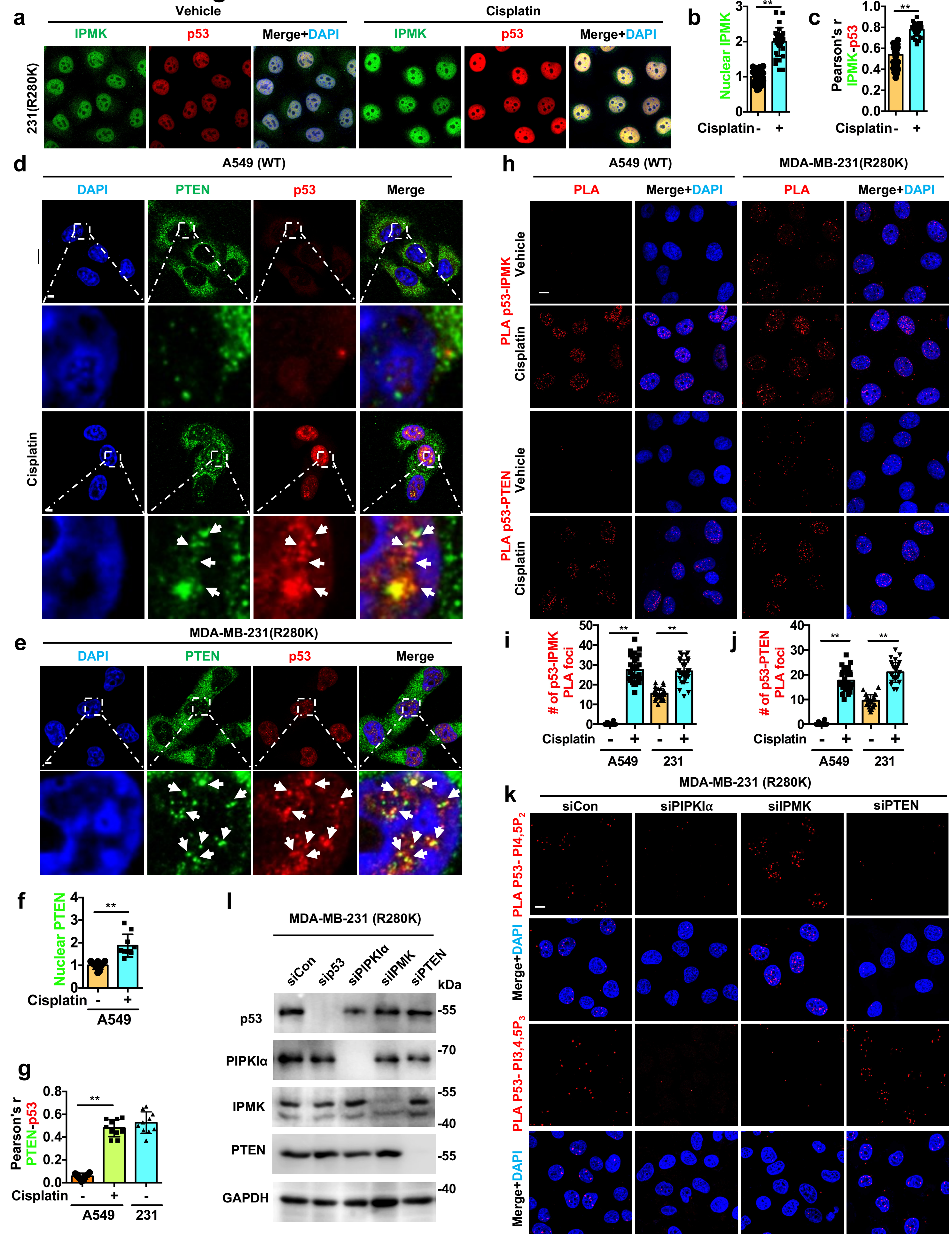

### Extended Data Fig. 6

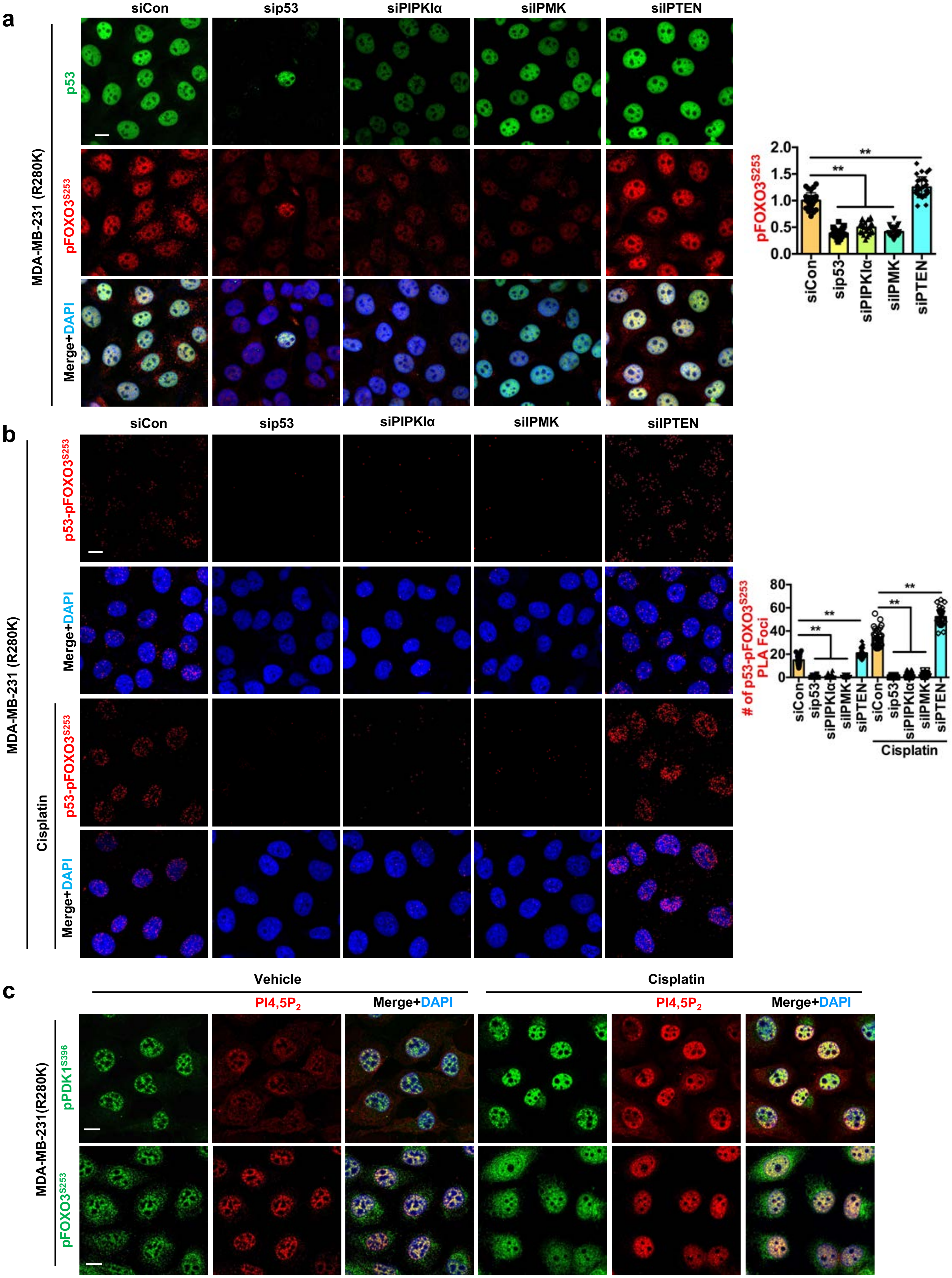

### Extended Data Fig. 7

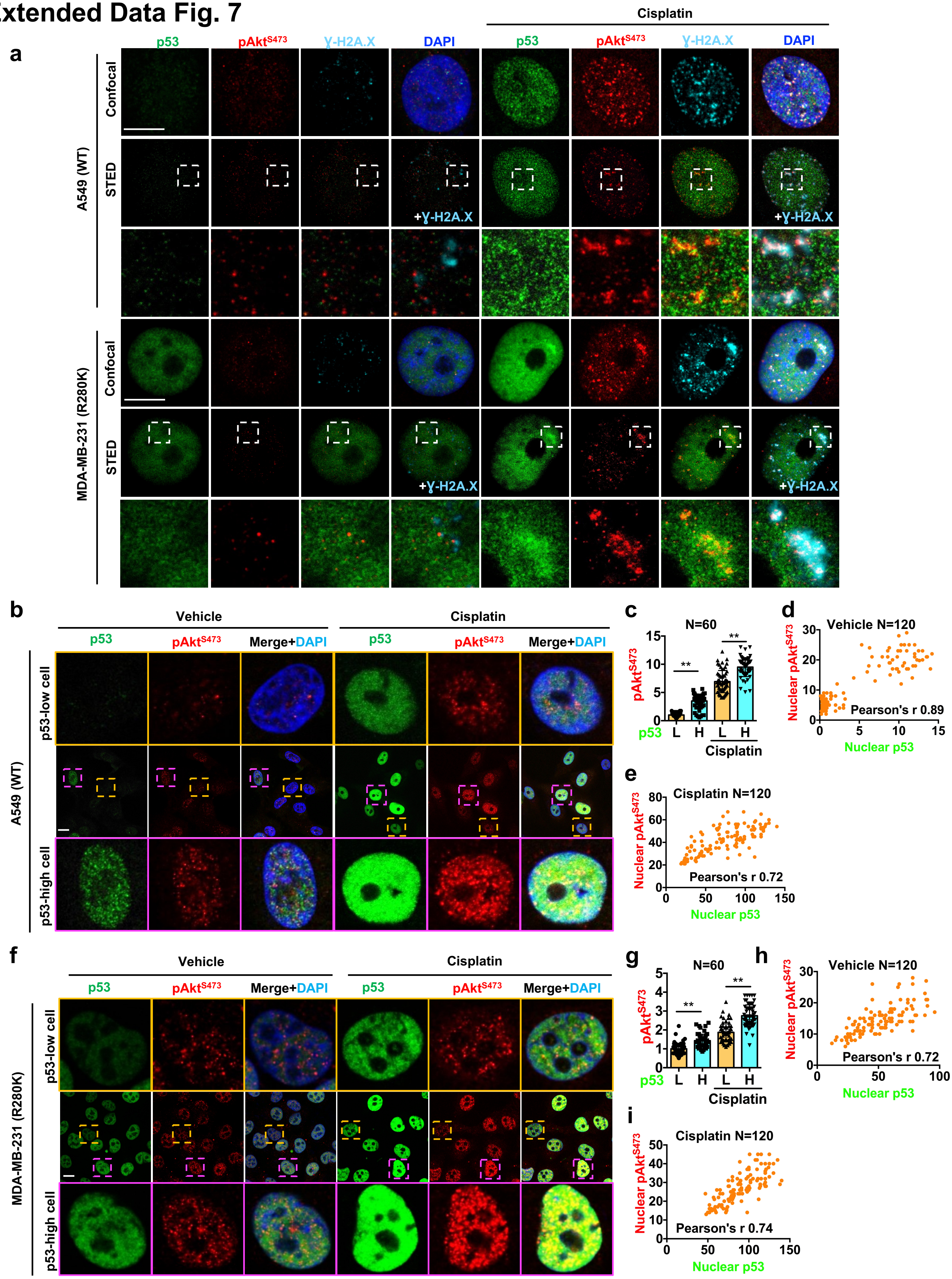

**Extended Data Table 1**

| Binding partners | Binding affinity with p53 (Kd) measured by MST |
|---|---|
| Akt1 | 10±2 nM |
| PDK1 | 6±0.2 nM |
| Sin1 | 20±1 nM |
| PIPKIα | 86±34 nM |
| IPMK | 42±13 nM |
| PTEN | 124±22 nM |
| PI (PolyPIPosome) | No binding detected |
| $PI4,5P_2$ (PolyPIPosome) | 150±12 nM |
| $PI3,4,5P_3$ (PolyPIPosome) | 1497±150 nM |
| PI (C16 Micelles) | No binding detected |
| $PI4,5P_2$ (C16 Micelles) | 83±7 nM |
| $PI3,4,5P_3$ (C16 Micelles) | 670±92 nM |
